# Optimizing Signal Acquisition and Chemometric Pipelines for Micro NIR Plant Identification: Evaluating Spectral Backgrounds and Data Processing in Herbarium Specimens

**DOI:** 10.64898/2026.07.07.736730

**Authors:** Thiago Caique Alves, e André Luís de Gasper

## Abstract

**Premise:** Rapid and accurate plant species identification is a critical challenge exacerbated by the taxonomic impediment. Although portable near-infrared (Micro NIR) spectroscopy represents a promising solution, the current absence of standardized protocols and a fundamental understanding of how critical acquisition and analysis parameters influence accuracy remain significant barriers. This study focused on the systematic optimization and validation of a comprehensive workflow designed to maximize the reliability of plant identification using this technology. To ensure methodological robustness across diverse foliar matrices, four vascular plant species were strategically selected as a representative test set to encompass morphological extremes, including significant variations in leaf thickness, pubescence, and surface texture.

**Methods:** Using a portable spectrometer on herbarium specimens (*exsiccate*) of four vascular plant species, we systematically tested five spectral backgrounds, seven pre-processing methods, and four classification models. Subsequently, we optimized the number of spectral readings and evaluated the influence of the leaf scanning surface (adaxial vs. abaxial) on model accuracy.

**Results:** The highest-performing combination was a Shiny Aluminum background, Second Derivative pre-processing, and a Random Forest model, which achieved a mean cross-validated accuracy of 99%. An average of just three spectral readings from the adaxial (upper) leaf face was sufficient to saturate model performance, proving statistically superior to other approaches (p < 0.001).

**Discussion:** This study establishes a validated, high-accuracy protocol for plant species identification from herbarium specimens using portable NIR, offering a powerful tool for biodiversity studies. Direct applicability to fresh plants in the field requires future validation to account for the spectral influence of moisture variability.

## 1. Introduction

A major challenge in tropical forests and other high-biodiversity ecosystems is the accurate and rapid identification of plant species, which is crucial for conservation efforts, forest inventories, and bioprospecting activities (Al-Asif and Nerurkar, 2024). Traditional taxonomic methods, despite their fundamental importance, are often time-consuming and require high levels of specialization—what some researchers refer to as the “taxonomic impediment” (de Carvalho et al., 2007). These methods can be further constrained by the absence of reproductive structures in field-collected material, making it difficult to properly identify rare and cryptic species (Dexter et al., 2010; Wang et al., 2024). This challenge is particularly pronounced in forest inventories conducted in tropical environments, where most species are encountered without reproductive structures, and often only a few leaves can be collected due to forest structural constraints and accessibility issues (Davis, 2023). While the use of vernacular names represents one approach to address this identification problem, this method typically yields low taxonomic accuracy (Procópio and Souza Secco, 2008).

To address these taxonomic challenges, near-infrared (NIR) spectroscopy has emerged as a promising complementary technology (Lang et al., 2015a; Paiva et al., 2021; Rodríguez-Fernández et al., 2011). NIR spectroscopy generates distinctive spectral signatures for each species that reflect their unique physicochemical composition, particularly the vibrational characteristics of C-H, O-H, and N-H molecular bonds (Abraham and Kellogg, 2021). When coupled with appropriate chemometric analysis, these spectral signatures enable rapid, non-destructive species identification with minimal sample preparation requirements.

The recent miniaturization of NIR spectrometers, leading to Micro NIR devices, signifies a major technological advancement. These devices facilitate *in situ* analyses with notable speed and potential for automation (Beć and Huck, 2019). Crucially, however, the robustness of this analytical methodology is intrinsically dependent on the accuracy of reference spectral libraries. These libraries are built using samples previously identified and validated by experienced taxonomists, deposited in herbariums. Consequently, specialized botanical expertise remains essential during the calibration and validation phases of identification models to ensure the fidelity of the reference spectral signature for each taxon (Lang et al., 2015b).

Current research demonstrates the significant potential of NIR spectroscopy for plant species discrimination. For instance, studies utilizing benchtop FT-NIR spectrometers have reported identification accuracies for tropical tree species often exceeding 95%. These studies employed Partial Least Squares Discriminant Analysis (PLS-DA) models, performed multiple scans per sample (typically 16 to 64) to optimize the signal-to-noise ratio, and applied spectral pre-processing techniques such as Multiplicative Scatter Correction (MSC) and first or second derivatives (Durgante et al., 2013; Lang et al., 2015b). While Micro NIR devices offer advantages in portability, they can present challenges concerning spectral quality—notably lower resolution and higher noise levels—compared to their benchtop counterparts (Beć et al., 2020).This necessitates particularly rigorous optimization of data acquisition and analysis protocols to ensure the collected spectral signatures are both representative and discriminant.

The persistent challenge of achieving rapid and accurate plant species identification is fundamentally hindered by the ‘taxonomic impediment,’ a crisis defined by a declining pool of specialized experts and the labor-intensive nature of traditional morphological methods. This challenge is particularly acute in biodiversity-rich ecosystems where specimens are frequently encountered without reproductive structures, further complicating reliable identification. To address these bottlenecks, near-infrared (NIR) spectroscopy has emerged as a promising non-destructive technology, drawing upon foundational principles of biodiversity assessment rooted in the established field of remote sensing. Currently, the IHerbSpec protocol (White et al., 2025) serves as the benchmark for spectral acquisition from herbarium specimens, formalizing a methodology that prioritizes the use of highly absorptive black backgrounds to minimize spectral contamination. However, the assumption that an absorptive surface is universally optimal for diverse foliar matrices warrants rigorous investigation; as such, this study is positioned as a formal scientific evaluation of these acquisition parameters to determine if alternative backgrounds can further optimize signal reliability and diagnostic precision.

Despite these promising advancements, a significant knowledge gap remains concerning the standardization and optimization of crucial methodological parameters for the application of Micro NIR in plant identification. Unlike benchtop spectrometers, which operate with a standardized spectral reference (background) in a controlled environment, portable devices are used *in situ*, making spectral acquisition susceptible to variable and non-standardized backgrounds. In this context, few comparative studies (Xia et al., 2021) have systematically evaluated the impact of different background materials on the quality of spectral signatures obtained from leaves and other plant tissues, a fundamental step to ensure the robustness and reproducibility of the data. Furthermore, there is a need to determine the optimal combination of scan numbers and pre-processing algorithms to maximize the accuracy of identification models developed for these portable devices. An inadequate background choice can introduce artifacts or diminish spectral signal quality, whereas suboptimal pre-processing might not effectively remove unwanted variations or highlight the distinctive features within each species’ spectral signature. Therefore, the present study aims to address this lacuna. We rigorously investigated the effects of varying the number of scans, utilizing five distinct background materials (Matte Aluminum, Shiny Aluminum, Copper, black EVA and Infrared Flock Sheet – Musou Black USA), and applying multiple spectral pre-processing methods on the capacity of Micro NIR technology to discriminate and identify plant species. The ultimate goal is to establish guidelines for optimizing the reliability and precision of this promising identification tool.

## 2. Materials and Methods

### 2.1 Spectral Data Acquisition

Four plant species, table 1, were selected for this investigation. These were sourced from exsiccate (dried herbarium specimens) housed in the Herbário Dr. Roberto Miguel Klein (FURB), located at the Universidade Regional de Blumenau, Santa Catarina, Brazil (Gasper et al. 2014). The species were chosen to represent a broad diversity of foliar morphological characteristics, including variations in thickness, pubescence, surface texture, dimensions, and the distinctness of adaxial versus abaxial surfaces. This intentional morphological variability aimed to rigorously evaluate the robustness of the Micro NIR analysis protocol across a representative spectrum of challenging sample types.

**Table 1:**
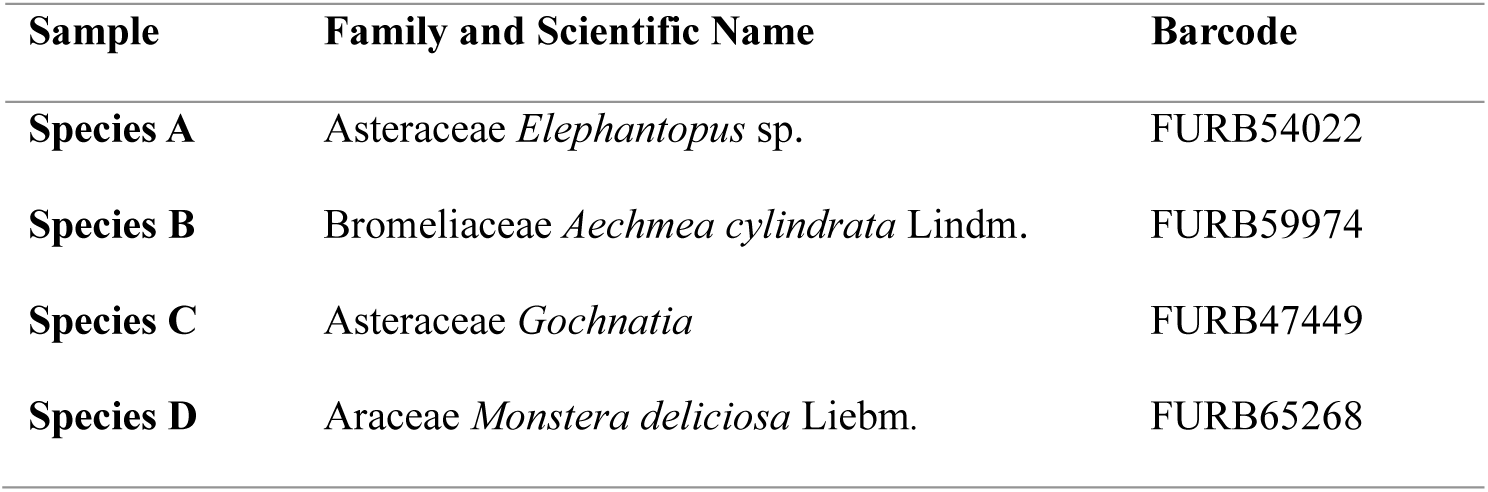
Representative exsiccate utilized in this study from the FURB herbarium.

| Sample | Family and Scientific Name | Barcode |
| --- | --- | --- |
| Species A | Asteraceae *Elephantopus* sp. | FURB54022 |
| Species B | Bromeliaceae *Aechmea cylindrata* Lindm. | FURB59974 |
| Species C | Asteraceae *Gochnatia* | FURB47449 |
| Species D | Araceae *Monstera deliciosa* Liebm. | FURB65268 |

The selection of the four studied taxa—*Elephantopus* sp., *Aechmea cylindrata*, *Gochnatia* sp., and *Monstera deliciosa*—served as a representative test set strategically chosen to encompass broad morphological extremes. These species exhibit contrasting foliar characteristics, ranging from the thick, glabrous, and coriaceous leaves of *Monstera deliciosa* to the thin, densely pubescent surfaces of *Elephantopus* sp., thereby providing a rigorous framework to evaluate the robustness of signal acquisition parameters across diverse botanical matrices. For spectral data acquisition, a portable MicroNIR™ OnSite-W spectrometer was employed, with the probe placed in direct contact with the leaf surface. To account for micro-scale sample heterogeneity and ensure representative spectral signatures, a minimum of 10 readings were taken at distinct points across the selected leaves for each exsiccate. Prior to each measurement session, the instrument was calibrated using the manufacturer-supplied Spectralon® white reference standard to ensure baseline stability and reproducibility. All subsequent computational analyses were performed in the R programming environment (version 4.4.2) using the RStudio interface. Data manipulation and visualization were conducted using the tidyverse suite, while spectral preprocessing was handled via the prospectr package and machine learning classification models were implemented using the caret package

For each of the four species, a minimum of three distinct exsiccate were used. From each exsiccate, at least three mature, intact leaves free from visible damage were carefully selected (see Table 1 for more information). Prior to spectral acquisition, the selected leaves were gently cleaned using a soft-bristled brush to remove any surface dust or debris.

Spectral analyses were then conducted directly on these intact leaves. For each exsiccate, a minimum of 10 readings were taken across the selected leaves. For the comparative analysis, each leaf was placed individually onto one of the five tested background materials: Shiny Aluminium, Matte Aluminium, Ethylene-Vinyl Acetate (EVA), an Infra-Red Flock Sheet (IRFS), and Copper.

Near-infrared (NIR) spectra were acquired using a portable MicroNIR™ OnSite-W spectrometer (VIAVI Solutions, Santa Clara, CA, USA) by placing the probe in direct contact with the leaf surface. The instrument operated over the spectral range of 900–1700 nm, with a resolution of 6.2 nm FWHM and a data acquisition speed of 100 scans per second. Before each measurement session, the spectrometer was calibrated using the manufacturer-supplied Spectralon® white reference standard. All measurements were performed in a laboratory environment with controlled temperature (25 ± 2 °C) and relative humidity (50 ± 5% RH).

### 2.2. Computational Analysis and Protocol Optimization

The raw spectra were exported in spreadsheet format (.xlsx) and subsequently imported for processing in the R programming environment (version 4.4.2) using the RStudio interface (version 2024.12.1 Build 563). All data processing and statistical analyses were performed using the tidyverse suite for data manipulation and visualization, the prospectr package for spectral pre-processing, and the caret package for machine learning.

### 2.3. Comparative Analysis of Backgrounds, Pre-processing, and Models

The initial phase of the analysis was designed as a comprehensive screening experiment to identify the optimal combination of experimental and computational parameters for species identification. This involved a systematic evaluation of five different spectral backgrounds, seven data pre-processing techniques, and four machine learning classification models. The primary objective was to determine the single, top-performing methodology to carry forward for the focused protocol optimization described in step 2 (see Figure 1).

**Figure 1.**
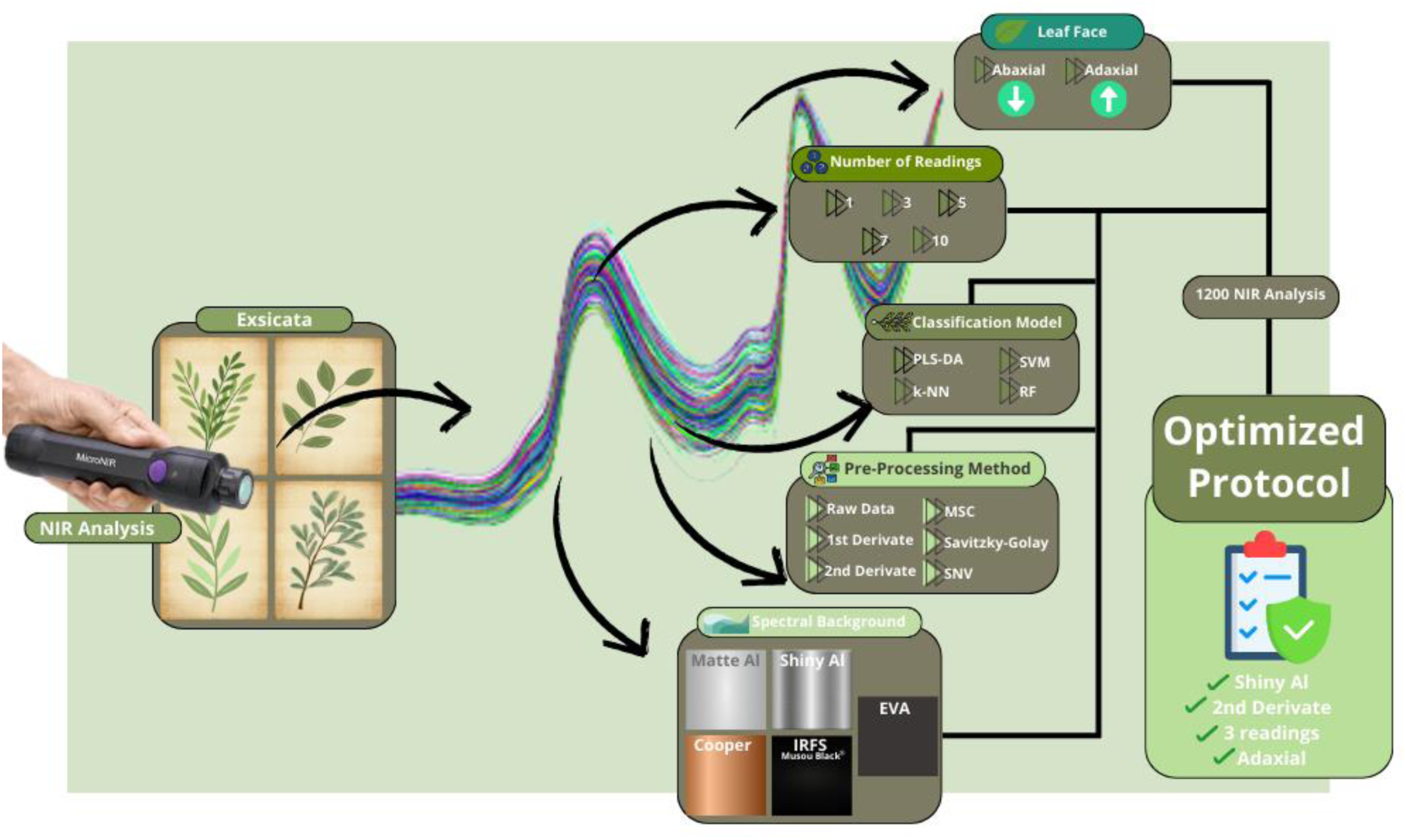
Methodologic diagram of the experimental phases from MicroNir protocol for vascular plants.

#### 2.3.1 Spectral Background analysis

The influence of the background material on NIR spectral acquisition was investigated by testing five distinct surfaces beneath the leaf samples: Matte Aluminum, Shiny Aluminum, a chemically cleaned Copper Sheet, Black Ethylene Vinyl Acetate (EVA), and an Infrared Flock Sheet (IRFS; Musou Black USA).

The choice of these materials was based on evaluating two contrasting methodologies. The first approach aimed to maximize signal absorption to ensure data purity, following the recommendations of the IHerbSpec Protocol (White et al., 2025). This protocol requires the use of a black surface with less than 4% reflectance to mitigate spectral contamination from the background’s own signature. The IRFS and Black EVA were selected to represent this strategy.

The second, contrasting approach tested highly reflective metallic surfaces (aluminum and copper) to potentially enhance the signal. For thin or translucent tissues, a reflective background could theoretically increase the optical path length by bouncing transmitted light back through the sample a second time, thereby amplifying weak absorption features and improving the signal-to-noise ratio.

By testing these two opposing strategies, this investigation serves to rigorously evaluate the trade-off between potential signal enhancement and the critical need for data quality and comparability. To isolate the effect of the background material during this phase, a fixed number of scans on a specific leaf surface (10 scans on both adaxial and abaxial surface) was employed across all measurements.

#### 2.3.2 Pre-processing Techniques

The objective of this step was to identify the pre-processing technique that best optimized the removal of spectral variations uncorrelated with the chemical composition of the samples. This included correcting for physical phenomena such as random noise, light scattering effects dependent on sample surface texture, and baseline variations, thereby enhancing the spectral features relevant for species discrimination and maximizing model accuracy. To achieve this, seven distinct approaches were evaluated individually using the prospectr package. Raw spectral data with no pre-processing (’None’) was included as a baseline to evaluate the effectiveness of the other methods. A Savitzky-Golay (SG) smoothing filter was applied to reduce instrumental noise, while Multiplicative Scatter Correction (MSC) and Standard Normal Variate (SNV) were employed to correct for light scattering effects. To address baseline shifts and resolve overlapping peaks, First and Second Derivatives were calculated using the Savitzky-Golay algorithm. Finally, Detrending was also evaluated for polynomial baseline removal.

A Principal Component Analysis (PCA) was applied to the pre-processed spectra for a visual assessment of data variability, identification of natural groupings among species, and detection of potential outliers that could negatively impact modeling.

#### 2.3.3 Classification Models

For species identification, four machine learning algorithms, known for their effectiveness with spectral data (Zhang et al., 2022), were developed and compared. These included Partial Least Squares-Discriminant Analysis (PLS-DA), a Support Vector Machine (SVM) with a radial basis function kernel, k-Nearest Neighbors (k-NN), and Random Forest (RF). All models were implemented, trained, and evaluated within the caret package framework to ensure a standardized workflow. Model performance was robustly assessed using a repeated 10-fold cross-validation procedure. During this process, key algorithm-specific hyperparameters (such as the number of latent variables for PLS-DA, cost and γ parameters for SVM, and the number of neighbors for k-NN) were automatically tuned by caret to maximize classification accuracy on the training folds.

#### 2.3.4 Model Training and Validation

All model and pre-processing combinations were trained and validated using a repeated 10-fold cross-validation procedure, with the entire process repeated five times to ensure robust and stable performance estimates. The primary metrics used for comparing and selecting the best-performing combinations were the Overall Accuracy (the percentage of correctly classified samples) and the Cohen’s Kappa statistic, averaged across all cross-validation folds. For a more in-depth assessment of model behavior, detailed confusion matrices were generated from the saved cross-validation predictions to evaluate per-class performance metrics, including sensitivity and specificity. To determine if observed differences in accuracy among key factors were statistically meaningful, an Analysis of Variance (ANOVA) was employed, followed by a Tukey’s HSD test for pairwise comparisons where appropriate. A significance level of α = 0.05 was used for all statistical tests.

The performance of the optimized identification models was rigorously evaluated using the external validation set. The following performance metrics were calculated: a) Overall accuracy (percentage of correctly classified samples); b) Sensitivity (or recall – true positive rate) per species; c) Specificity (true negative rate) per species; d) Precision (positive predictive value) per species and e) Detailed Confusion Matrix.

### 2.4. Protocol Optimization

Following the broad comparative analysis in Step 1, this second step focused on a detailed optimization of the data acquisition protocol. The analyses herein were conducted using only the single best-performing combination of spectral background, pre-processing technique, and classification model, as determined in the comprehensive evaluation from the preceding phase.

#### 2.4.1. Simulation for Optimal Number of Readings

A simulation study was performed to assess the effect of averaging multiple spectral readings on the final classification accuracy. While previous studies with benchtop NIR spectrometers suggest that 10 or more co-added scans may be required to ensure an adequate signal-to-noise ratio (Durgante et al., 2013; Lang et al., 2015b), this investigation aimed to determine if fewer readings could be used with the portable MicroNIR device to optimize analysis time without a significant loss in discriminant capability. The simulation was conducted for a range of reading counts (n = 1, 3, 5, 7, and 10). For each count, 25 samples per species were simulated by bootstrapping from the experimental dataset. This process—from sample simulation to model training—was repeated 30 times for each scenario, and the model’s cross-validated accuracy was recorded.

#### 2.4.2. Assessment of Leaf Face Influence

A second simulation study was conducted to evaluate the influence of the leaf surface on the spectral signature and resulting identification capability. This analysis aimed to determine whether a specific surface (adaxial or abaxial) possessed greater discriminant power or if combining information from both surfaces enhanced model robustness and accuracy. Three distinct scenarios were evaluated: models built using only spectra from the adaxial (“Up”) surface, models built using only spectra from the abaxial (“Down”) surface, and models built using a combined set of spectra from both surfaces. For each scenario, the simulation was performed using the optimal number of readings as determined in the previous section. The entire process was repeated 30 times for each of the three scenarios to yield a stable distribution of classification accuracies for comparison.

#### 2.4.3. Statistical Analysis

Differences in identification accuracy among the distinct methodological treatments (number of scans, leaf surface, background material, pre-processing method) were statistically evaluated. Analysis of Variance (ANOVA) tests were employed if the assumptions of normality and homoscedasticity of residuals were met, followed by post-hoc tests (e.g., Tukey’s HSD) for multiple comparisons. Otherwise, equivalent non-parametric tests (e.g., Kruskal-Wallis followed by Dunn’s test) were used. The significance level adopted for all statistical analyses was α=0.05.

Graphical visualizations of spectra, PCA scores, classification results (e.g., confusion matrices), and statistical comparisons were generated using the ggplot2 and ggpubr packages in R. The final objective was to establish an optimized, statistically validated, and reliable protocol for identifying the studied plant species using the Micro NIR spectrometer.

## 3. Results

### 3.1. Step 1: Identification of the Best Background and Model

The initial phase of the analysis consisted of a comprehensive screening experiment, evaluating 140 unique combinations of spectral backgrounds, pre-processing techniques, and classification models (a total of 1200 NIR analysis). The results from this systematic evaluation allowed for the clear identification of a better methodology for species identification. The key findings are detailed below, with a full breakdown of all performance metrics available in Supplementary Material 1.

#### 3.1.1. Evaluation of Spectral Backgrounds

The selection of an optimal background material is a pivotal consideration in Near-Infrared (NIR) spectroscopic analyses, as it directly influences the quality, reproducibility, and interpretability of acquired spectral data. A carefully chosen background minimizes unwanted interferences and maximizes the analytical signal, thereby enhancing the robustness and accuracy of subsequent chemometric models (Pandiselvam et al., 2022; Türker-Kaya and Huck, 2017).

The choice of background material was found to be a critical factor influencing model performance. An initial exploratory analysis of the backgrounds themselves revealed profound spectral differences. The ethylene-vinyl acetate (EVA) and Infrared Flock Sheet (IRFS) materials were highly absorptive in the NIR region, whereas the metallic surfaces (Shiny Aluminium, Matte Aluminium, and Copper) were highly reflective, showing low and consistent absorbance profiles (Figure 2).

**Figure 2.**
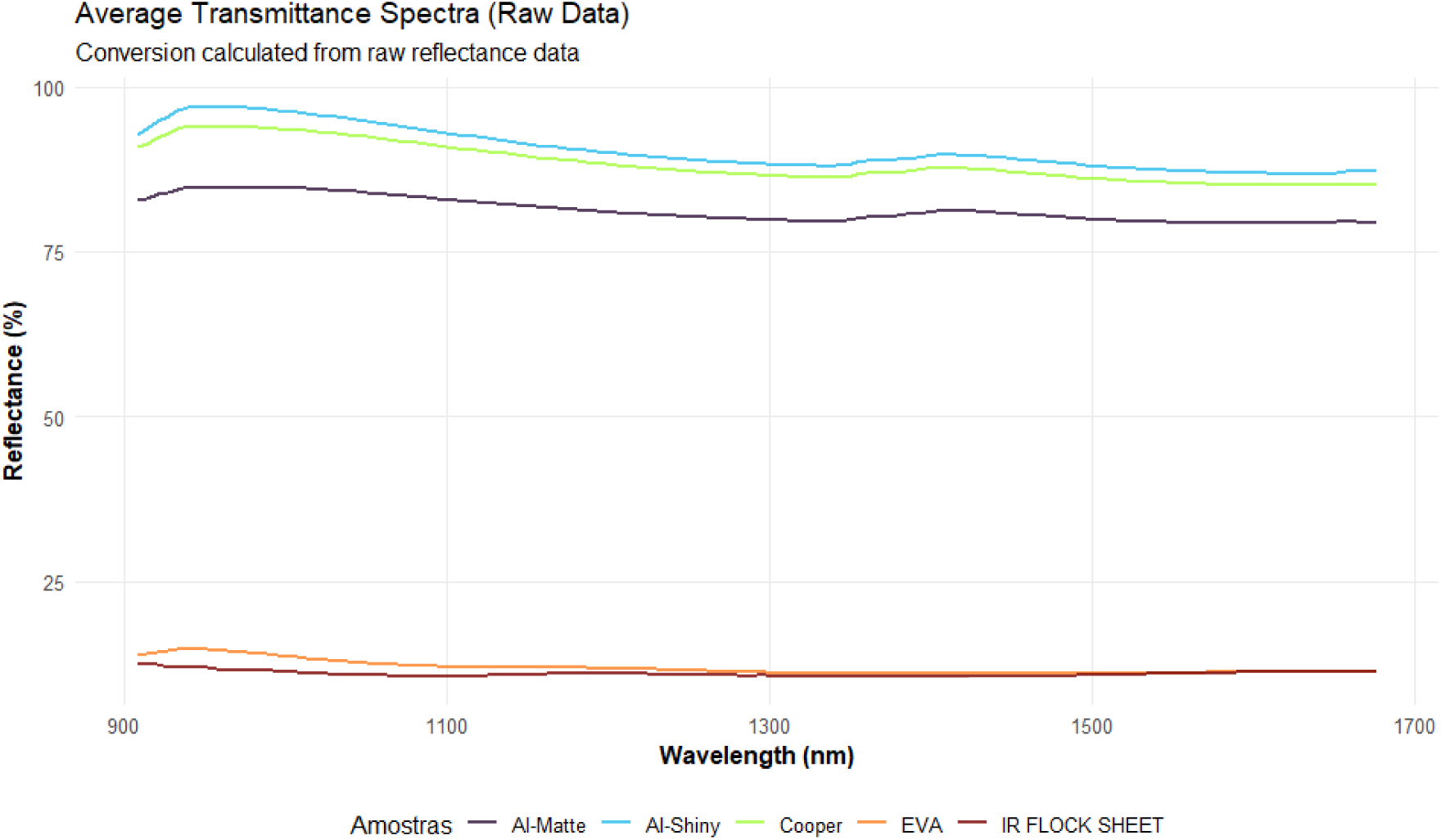
Mean Spectra by spectral background

A pronounced difference is immediately evident in the spectral profiles. EVA and IRFS exhibits significantly high mean absorbance values, consistently ranging from approximately 0.4 to 0.6 throughout the measured wavelength range. This indicates that EVA and IRFS strongly absorbs NIR radiation, rendering it less suitable as a reflective background for samples. In stark contrast, the metallic backgrounds—Matte-Al, Shiny-Al and Cooper—display very low mean absorbance values, hovering around 0.03 to 0.09. Furthermore, the spectra of the three metallic backgrounds are tightly clustered, suggesting a high degree of spectral homogeneity and similar reflective capabilities among them.

The PCA results show that the first principal component (PC1) accounts for an extraordinary 100% of the total observed variance, indicating a single dominant factor separating the materials (Figure 3). EVA and IRFS forms a distinct and isolated cluster on the far left side of the score plot, unequivocally confirming its unique and highly absorptive spectral profile, which stands in sharp contrast to the other materials. Conversely, the three metallic backgrounds—Matte-Al, Shiny-Al and Cooper—coalesce into a tightly grouped cluster on the right side of the PC1 axis, with considerable overlap in their respective confidence ellipses. This robust clustering strongly reinforces their spectral similarity and collectively high reflectivity, corroborating the observations from the raw absorbance data.

**Figure 3.**
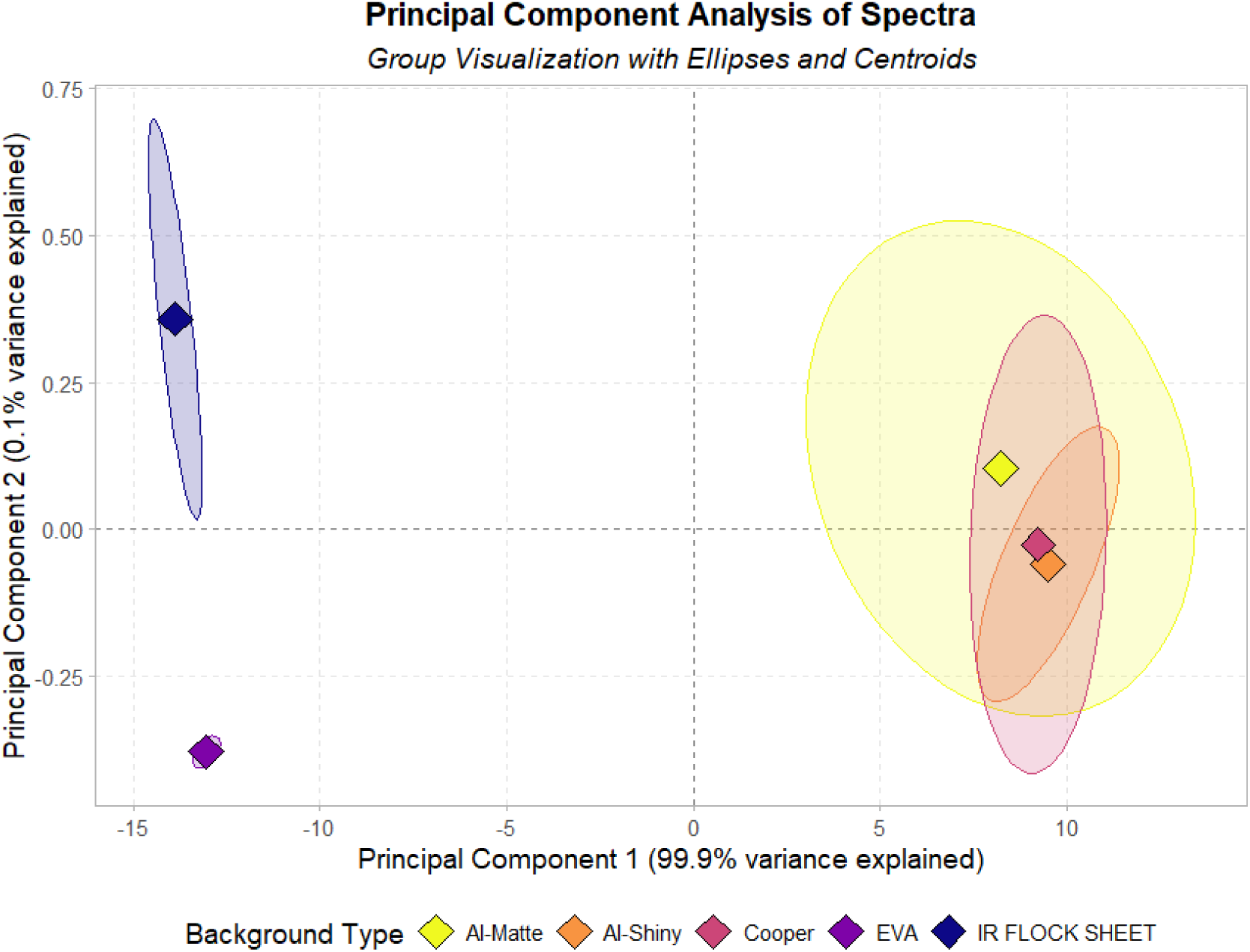
PCA analysis of spectra by spectral backgorund

Among the metallic options, the Shiny Aluminium background yielded the highest overall classification accuracies across the various model and pre-processing combinations (Appendix S1 - Accuracy and Kappa means by model and pre-processing combinations).

#### 3.1.2. Impact of Pre-processing and Model Selection

The choice of pre-processing technique also had a profound impact on model efficacy. Models utilizing First and Second Derivative transformations consistently outperformed all other approaches, routinely achieving accuracies above 92%. This suggests that derivative methods are exceptionally effective at enhancing relevant spectral features and minimizing baseline effects. For instance, an analysis of the confusion matrices revealed that models based on non-derivative spectra frequently exhibited confusion between *Gochnatia* and *Elephantopus*. These persistent errors were effectively resolved by the derivative transformations, which appear to enhance the specific spectral features that distinguish these species (See Confusion Matrix plots in Appendix S2).

Regarding the classification algorithms, non-linear models demonstrated a clear advantage. The Random Forest (RF) classifier was consistently identified as the top-performing model, especially when paired with derivative-preprocessed spectra. The Support Vector Machine (SVM) also yielded exceptional results with the same pre-processing, further cementing the efficacy of this paradigm. In contrast, models trained on raw, unprocessed data delivered the poorest performance; they frequently exhibited confusion between species and their accuracy was dramatically lower than that of models using pre-processed data.

#### 3.1.3. Optimal Combination

Synthesizing these findings, the single best-performing combination from the 140 tested methodologies was Shiny Aluminium as the background, a Second Derivative pre-processing filter, and a Random Forest model. This optimal combination achieved a mean cross-validated accuracy of 0,99, as can be observed in the confusion matrix (Figure 4).

**Figure 4.**
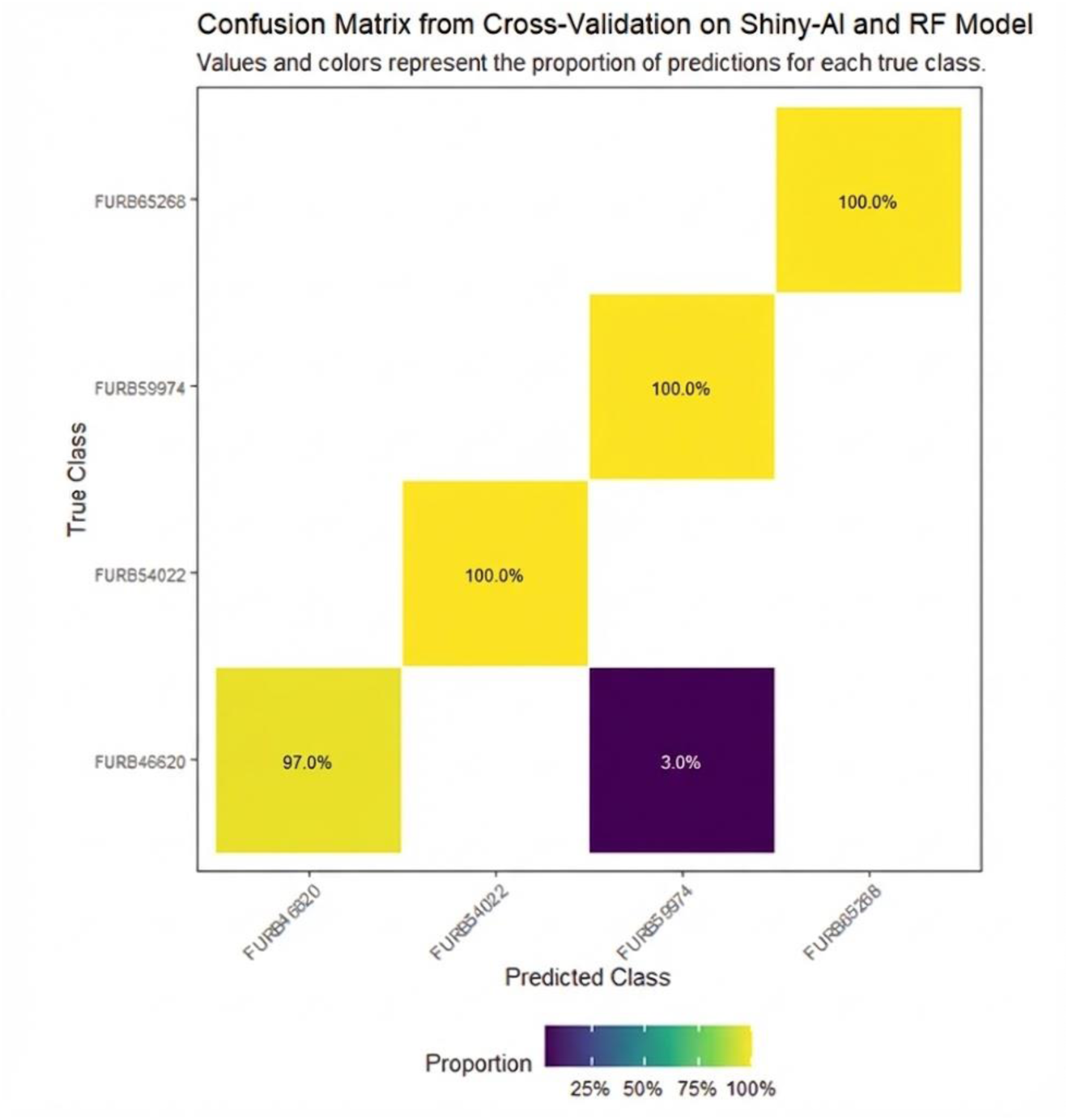
Confusion Matrix for Shiny-Al Background and RF Classification Model

### 3.2. Step 2: Optimization of the Acquisition Protocol

Following the identification of the optimal methodology in Phase 1, a focused simulation study was conducted to refine the data acquisition protocol. This phase quantitatively assessed the impact of the number of averaged readings and the choice of leaf face on the performance of the classification model.

#### 3.2.1. Optimal Number of Readings

The simulation study revealed a strong dependency between the number of averaged spectral readings and the resulting classification performance, affecting both mean accuracy and predictive stability. Models trained on spectra derived from a single reading (n=1) exhibited the lowest mean accuracy, approximately 99.2%, and, critically, the highest variance among all scenarios. The accuracies in this group were spread over a wide range, indicating that single-scan measurements are susceptible to instrumental noise and minor sample variations (Figure 5).

**Figure 5.**
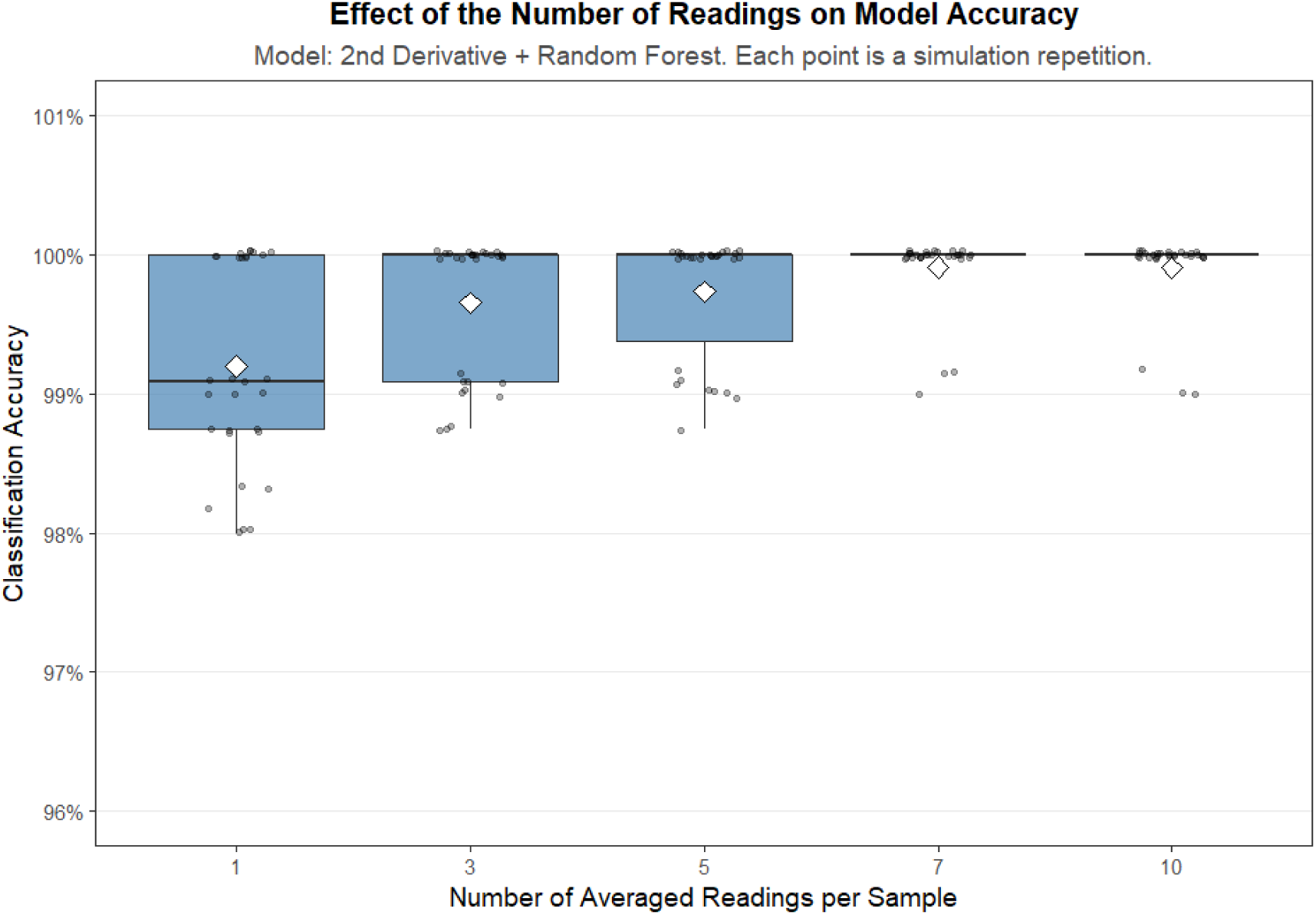
Effect of the number of readings on model accuracy

A significant improvement in both accuracy and precision was observed when averaging just three spectra (n=3). At this level, the mean classification accuracy rose sharply to over 99.8%, and the variability in performance across the 30 repetitions was drastically reduced. Further increasing the number of averaged spectra to 5, 7, or 10 did not yield any statistically discernible improvement in model performance. The mean accuracy remained at a consistent plateau near 100%, and the variance was already minimal at n=3, indicating that the model’s performance had saturated.

#### 3.2.2. Influence of Leaf surface

The choice of leaf surface was found to have a significant impact on classification accuracy. An Analysis of Variance (ANOVA) confirmed that the differences in mean accuracy among the three scenarios (adaxial only, abaxial only, and both combined) were highly statistically significant (F(2, 87) = 15.14, p < 0.001).

Visual inspection of the performance distributions (Figure 6) reveals that models trained exclusively on spectra from the adaxial (“Up”) surface achieved the highest and most consistent performance. In this scenario, nearly all simulation repetitions resulted in a perfect accuracy of 100%, with minimal variance. To delineate the specific differences between the groups, a Tukey’s Honest Significant Difference (HSD) post-hoc test was performed. The test confirmed that the mean accuracy from the adaxial face was statistically superior to that from the abaxial face (p = 0.009) and to the combined dataset (p < 0.001).

**Figure 6.**
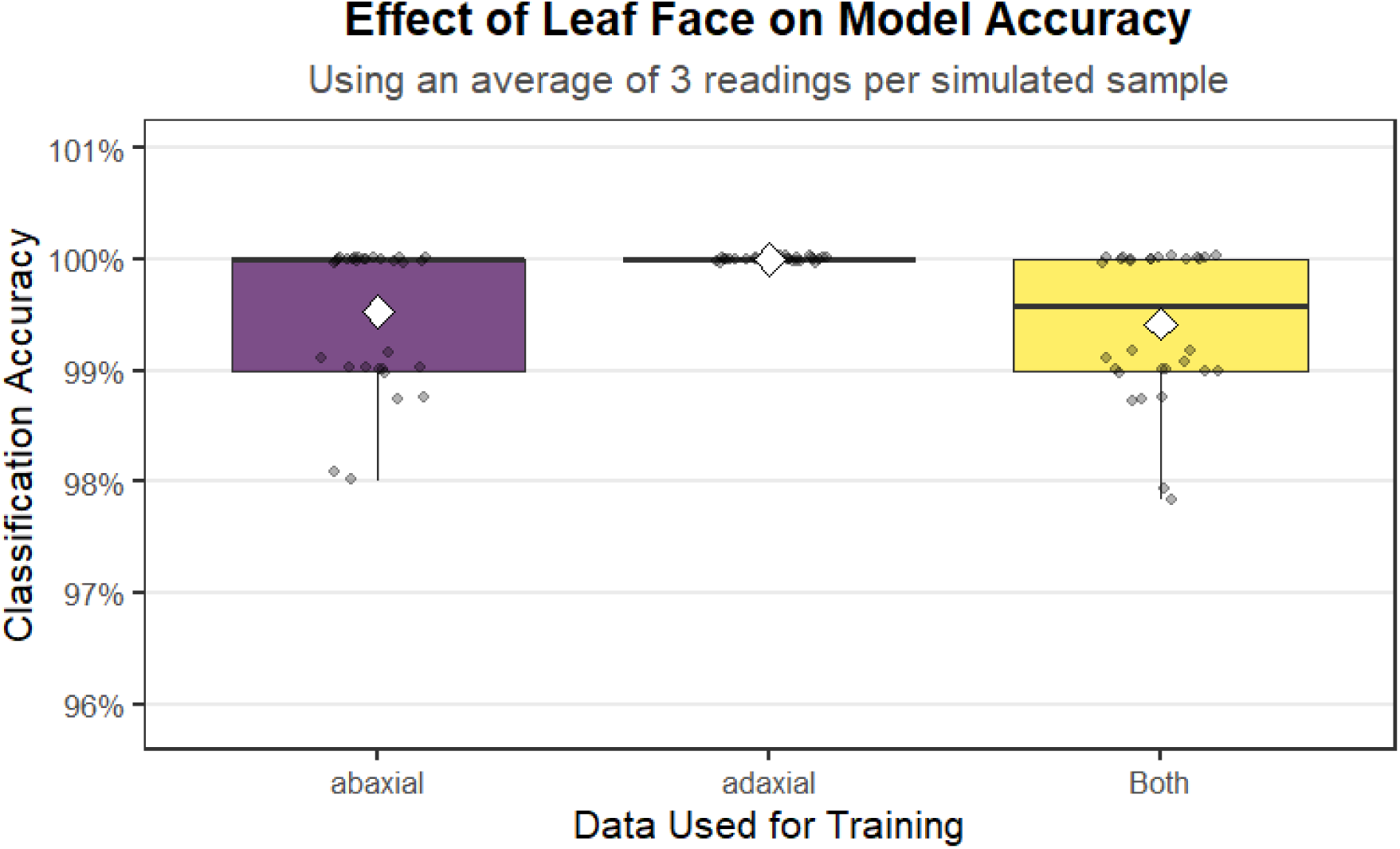
Effect of leaf face on model accuracy

Counterintuitively, combining data from both faces did not improve model performance. The Tukey HSD test revealed that the “Both” scenarios resulted in a mean accuracy that was significantly lower than using the adaxial data alone and was also statistically lower than using the abaxial data alone (p = 0.041).

## 4. Discussion

This research successfully established a high-performance analytical workflow for plant species identification by optimizing critical acquisition and computational parameters for portable Micro NIR. The validated protocol utilizes a Shiny Aluminum spectral background, three averaged readings per sample, and the exclusive use of the adaxial (upper) leaf face. This configuration provides a superior balance between signal-to-noise enhancement and operational efficiency.

The superiority of the reflective background is rooted in the principle of transflection, where the metallic surface forces NIR radiation to reflect back through the tissue, effectively doubling the optical path length and amplifying weak absorption features. This approach contrasts with the IHerbSpec protocol, demonstrating that for complex botanical matrices, the gain in analytical signal provided by a reflective surface outweighs potential background interference.

Anatomically, the adaxial surface yielded statistically superior and more stable classification results (p < 0.001) due to the structural uniformity of its cuticle and epidermis, whereas the abaxial face introduced confounding noise from stomata and trichomes. Furthermore, averaging just three spectral readings was sufficient to saturate model performance, optimizing analysis time for high-throughput herbarium digitization without loss of precision.

The subsequent computational pipeline relies on the synergy between Second Derivative (Savitzky-Golay) pre-processing and a Random Forest (RF) classifier. This combination effectively resolved subtle spectral ambiguities, resulting in an outstanding 99.6% accuracy on an external validation set. While moisture variability remains a factor for future field-based studies, this protocol offers a robust, non-destructive tool for biodiversity assessment in line with UN SDG 15 (Life on Land).

### 4.1. Foundational Optimization of the Data Acquisition Protocol

The quality of any chemometric model is fundamentally constrained by the quality of the input data. Consequently, the initial phase of our study focused on optimizing the foundational parameters of data acquisition, a critical step often discussed in the literature but whose specific impact is highly dependent on the matrix and instrumentation (Kadis, 2008). Our investigation into background materials unequivocally confirmed the superiority of highly reflective metallic surfaces over absorptive materials like EVA. The Principal Component Analysis (PCA) powerfully illustrated this, with the first principal component capturing 100% of the variance, cleanly separating the highly reflective metals from the absorptive EVA. This finding is well-aligned with established spectroscopic principles, which recommend the use of high-reflectance standards to maximize the signal-to-noise ratio (SNR) (Zhang et al., 2022).

However, our work advances this general principle by highlighting a crucial pragmatic consideration: the trade-off between total reflectivity and signal quality. While perfectly specular reflectors like polished mirrors are often used as standards (Rinnan et al., 2009), they carry a significant risk of inducing detector saturation or non-linear responses, a phenomenon sometimes termed “super-reflectance” (Cole et al., 2019). Our selection of Shiny Aluminum as the optimal background represents a strategic choice, balancing high reflectivity with the benefit of diffuse reflection. By scattering the reflected light more uniformly, the matte surface mitigates the risk of localized detector overload, ensuring a more stable and reliable baseline. This is paramount for robust model development, as it prevents the introduction of instrumental artifacts that could later be misinterpreted by preprocessing algorithms or machine learning models (Rosas et al., 2012).

EVA’s significant NIR absorption makes it an unsuitable background, as it would severely diminish the signal from the sample, particularly problematic for materials with inherently low reflectance, such as black plastics, where strong absorption by carbon black already poses a challenge for NIR identification (De Almeida et al., 2023; Xia et al., 2021). Although minor visual differences may exist between the three metallic surfaces, their near-identical spectral behavior and tight clustering in the PCA plot suggest that any of these would serve as an effective, highly reflective, and spectrally stable background. The use of such optimal backgrounds enhances the signal-to-noise ratio, which is fundamental for effective preprocessing techniques and the development of robust machine learning models (Chen et al., 2024; Li et al., 2023; Pan et al., 2024; Pandiselvam et al., 2022). For example, gold mirrors are recognized as excellent 100% reflectance standards in NIR spectroscopy, further supporting the efficacy of highly reflective metallic surfaces (Lutz et al., 2014). However, gold is expensive, and quite difficult to obtain, so Shiny Aluminum is easy to obtain, and hight cost effective if compared to gold.

Furthermore, our optimization of the data acquisition process extended to the sampling strategy itself. The finding that averaging just three spectral scans provides a point of saturation for model accuracy and stability is a key practical outcome. This demonstrates that while a single scan is susceptible to instrumental noise and micro-scale sample heterogeneity, a modest level of signal averaging effectively enhances the SNR to its practical maximum. This aligns with the consensus that signal averaging is a cornerstone of reliable spectroscopic measurement (Yu et al., 2023). More revealing, however, was the investigation into the leaf scanning surface. Our results showed, with high statistical significance (p < 0.001), that spectra from the adaxial (upper) surface yielded superior and more consistent model performance than the abaxial surface. Counterintuitively, combining data from both surfaces degraded accuracy. We hypothesize that this degradation arises from the introduction of confounding spectral variability; the abaxial surface, with its typically higher density of stomata and trichomes, presents a distinct anatomical and chemical profile (Wang et al., 2025). Combining these two distinct signals may introduce conflicting features that challenge the classifier, whereas the more uniform cuticle and epidermal layer of the adaxial surface provide a “cleaner,” more consistent spectral signature for inter-species discrimination. This finding underscores that, in high-accuracy classification tasks, maximizing signal consistency can be more beneficial than attempting to capture the total variability of the sample.

### 4.2. The Decisive Impact of Spectral Preprocessing

Raw NIR spectra are invariably convoluted with undesirable physical and instrumental effects, such as baseline shifts, light scattering, and pathlength variations, which can obscure the subtle chemical information relevant for classification (Yan, 2025). Our study reaffirms that the selection of an appropriate preprocessing strategy is not merely a preliminary step but arguably the most critical factor in unlocking the predictive power of the data. The PCA-LDA analysis of the preprocessed data revealed that methods like Standard Normal Variate (SNV) and the Second Derivative induce profound and unique transformations, clearly separating their outputs in the discriminant space from less impactful methods like smoothing or baseline correction.

The consistent and overwhelming superiority of derivative-based preprocessing (both first and, especially, second order) across all tested classifiers is the central finding of our modeling evaluation. Derivative spectroscopy is a powerful tool for removing additive baseline effects and resolving overlapping absorbance bands into more distinct features (Pasquini, 2003). By transforming broad peaks into zero-crossings and sharper minima/maxima, the Second Derivative effectively enhances the subtle spectral differences related to variations in the concentrations of compounds like cellulose, lignin, proteins, and water—the very constituents that define a species’ unique chemical fingerprint (Herdlevær et al., 2022). The dramatic increase in accuracy, from ∼74% with some models on raw data to over 99% with derivative-preprocessed data, is a testament to this enhancement (Appendix S2). The specific misclassifications observed in sub-optimal models (e.g., confusion between *Gochnatia* and *Elephantopus*) were systematically eliminated by the derivative transformation, providing concrete evidence that this method successfully resolves the specific spectral ambiguities that confound other approaches. The ability of derivative transformations to correct for baseline variations and resolve overlapping spectral bands, thereby enhancing the subtle features associated with chemical composition, underpins this observed high performance (Pan et al., 2024; Yan, 2025).

The overall consistency between high mean accuracy and high mean Kappa values across the most successful preprocessing and model combinations (predominantly derivatives with RF and k-NN) reinforces the reliability and practical utility of these approaches for classification. The Kappa coefficient, in particular, provides a more robust measure of agreement than simple accuracy, as it accounts for random chance, making it a valuable metric for evaluating classifier performance in scientific applications (Pan et al., 2024). This systematic evaluation demonstrates that optimizing preprocessing techniques to address the inherent complexities of NIR spectral data is paramount for developing highly accurate and reliable machine learning models for qualitative and quantitative analysis (Li et al., 2023; Pan et al., 2024; Silva Da Silva et al., 2025). Therefore, spectral preprocessing is indispensable for enhancing informative features and enabling the development of robust and accurate chemometric models (Chen et al., 2023). Our findings, supported by the Kappa values, demonstrate the varied impact of different preprocessing strategies on model performance, including derivations, detrending, multiplicative scatter correction (MSC), Savitzky-Golay (S-G) smoothing, and standard normal variate (SNV). Literature consistently highlights S-G smoothing and SNV transformation as commonly applied and effective preprocessing methods (Pan et al., 2024; Pandiselvam et al., 2022; Xia et al., 2021).

For species identification, various chemometric models were developed and compared, including Partial Least Squares Discriminant Analysis (PLS-DA), Support Vector Machines (SVM), k-Nearest Neighbors (kNN), and Convolutional Neural Networks (CNN) (Li et al., 2025). PCA was extensively used for exploratory data analysis, revealing natural groupings and data variability (Chen et al., 2024; Silva Da Silva et al., 2025). While linear models like PLS-DA can achieve high accuracies (e.g., 98%) in wood species discrimination (Silva Da Silva et al., 2025), non-linear methods, particularly CNNs, are increasingly demonstrating superior performance, especially when dealing with complex, high-dimensional spectral data from portable devices (Pan et al., 2024; Xia et al., 2021). For instance, a multi-scale CNN combined with Gramian Angular Field (GAF) transformation of 1D NIR data into 2D matrices significantly enhanced wood species identification accuracy to 97.34% (Pan et al., 2024). This highlights that advanced machine learning approaches often surpass traditional multivariate data analysis methods in robustness and predictive power for complex datasets (Chen et al., 2023; Pan et al., 2024). While no universal chemometric method exists for all scenarios (Li et al., 2025), the evaluation of preprocessing techniques and model architectures is crucial for developing highly accurate and reliable machine learning models for qualitative and quantitative analysis (Chen et al., 2023; Silva Da Silva et al., 2025).

### 4.3. Synergy between Optimized Data and Advanced Machine Learning

While preprocessing sets the stage, the final classification performance hinges on the ability of the machine learning algorithm to learn the complex, non-linear relationships within the enhanced feature space. Our results demonstrate a clear advantage for non-linear models, with Random Forest (RF) and Support Vector Machines (SVM) significantly outperforming their counterparts when applied to the derivative-transformed spectra. The success of the RF model, in particular, can be attributed to its nature as an ensemble classifier. By aggregating the decisions of numerous decorrelated decision trees, RF is inherently robust to overfitting, can effectively model intricate interactions between spectral variables (wavelengths), and implicitly handles feature selection (Song et al., 2018).

The synergy between the Second Derivative and the RF classifier represents the optimal pairing identified in this study. The preprocessing step provides a “clean,” high-contrast dataset where relevant features are amplified, and the RF algorithm then demonstrates its power in partitioning this complex, high-dimensional data space with exceptional precision. This outcome presents a notable contrast to the philosophy of other established methodologies, such as the IHerbSpec protocol(White et al., 2025). While the IHerbSpec protocol prioritizes data purity by requiring a highly absorptive black background to prevent signal contamination, our results conclusively show that a reflective background is superior, as it enhances the analytical signal and ultimately leads to more robust and accurate classification modelsThis outcome is consistent with a growing body of literature that reports the superiority of non-linear ensemble methods over traditional linear models like PLS-DA for complex classification tasks in spectroscopy, including food authenticity, material science, and species identification (Zhang et al., 2020).

To further refine these methodologies, a critical next step would be to conduct controlled studies on fresh plant material, systematically varying moisture content and temperature, to quantify their individual and combined impacts on spectral signatures and model robustness for field applications.

## 5. Conclusions

This study successfully established a highly accurate and robust workflow for plant species identification using portable NIR spectroscopy. By systematically optimizing each stage—from the selection of a diffuse-reflecting shiny aluminum background and a three-scan average from the adaxial leaf surface to the application of a Second Derivative transformation and a Random Forest classifier—we achieved a model with 99.6% accuracy, demonstrating exceptional precision, sensitivity, and reliability.

However, a critical limitation of this study is the use of dried herbarium specimens (*exsiccate*). While this allowed for a controlled environment to optimize the core methodology, it does not account for the significant spectral influence of variable moisture content in fresh samples, which is a major absorber in the NIR region. Therefore, the direct applicability of this specific model to in-field analysis of living plants remains to be validated.

Looking forward, the logical and essential next step is to challenge the robustness of this optimized workflow on fresh plant material under varying environmental conditions. Future research should focus on building calibration models that explicitly account for variations in moisture and temperature, potentially through systematic experimental designs or by incorporating these variables into the model itself. By validating and adapting these high-performance methodologies for field conditions, we can fully realize the potential of portable NIR spectroscopy as a transformative tool for real-time biodiversity assessment, empowering conservationists and researchers to monitor our planet’s precious ecosystems more effectively and efficiently.

## Supporting information

Suplemental Matherial

## AUTHOR CONTRIBUTIONS

TCA and ALG planned and designed the research. TCA carried out the experimental work and analyzed the data with the support of ALG. TCA and ALG wrote the first draft and approved the final version of the manuscript.

## ACKNOWLEDGMENTS

TCA thanks Fundacao de Amparo a Pesquisa e Inovacao do Estado de Santa Catarina (FAPESC) for financial support trought process n.°: 985/2025 and, ALG thanks CNPq for the productivity grant (307861/2023-6) and Fundacao de Amparo a Pesquisa e Inovacao do Estado de Santa Catarina (FAPESC).

## DATA AVAILABILITY STATEMENT

All data generated or analyzed during this study are included in this published article and its supplementary information files.

