## Supplementary material for "Optimizing Signal Acquisition and Chemometric Pipelines for Micro NIR Plant Identification: Evaluating Spectral Backgrounds and Data Processing in Herbarium Specimens": Suplemental Matherial

Supplementary Material: Analysis of FTIR Spectra on Different Backgrounds


#### Table of contents

- Introduction
- 1. Setup: Loading Required Packages
  - 2. EVA Background
    - 2.1. Data Loading and Preparation
    - 2.2. Model Training Pipeline Setup
    - 2.3. Executing the Training and Evaluation Loop
    - 2.4. Analysis and Visualization of Results
    - **2.5 Comprehensive Pipeline for Pre-processing and Model Evaluation**
    - **2.6 Graphical Analysis of Cross-Validation Confusion Matrices**
    - **2.7 Visualization of Mean Spectra by Pre-processing Method**
- 3. Shiny Alluminiun Background

### Supplementary Material: Analysis of FTIR Spectra on Different Backgrounds

 Code

- Show All Code
- Hide All Code
- ---
- View Source

### Introduction

This document provides the supplementary material for our study on plant species identification using Fourier-transform infrared (FTIR) spectroscopy. Here, we detail the computational methodology used to test and compare the performance of various machine learning models combined with different spectral pre-processing techniques.

### 1. Setup: Loading Required Packages

First, we load all the necessary R packages for data manipulation, pre-processing, modeling, and visualization. A brief description of each package’s role is provided in the comments.

Code

```
# Data Manipulation and Visualization
library(readxl)     # To read data from Excel files
library(tidyverse)  # A collection of R packages for data science (incl. dplyr, ggplot2)
library(ggplot2)    # For creating elegant and complex plots
library(ggpubr)     # For arranging and customizing ggplot2 plots
library(viridis)    # For colorblind-friendly color palettes

# Spectral Data Pre-processing
library(prospectr)  # Provides a wide range of chemometric tools, including pre-processing
library(signal)     # Used for signal processing functions like Savitzky-Golay filter

# Machine Learning and Model Evaluation
library(caret)      # Comprehensive framework for training and evaluating machine learning models
library(pls)        # Implements Partial Least Squares (used for PLS-DA)
library(randomForest) # Implements Random Forest algorithm
library(kernlab)    # Implements Support Vector Machines
library(e1071)      # General-purpose package with various ML functions

# Reporting
library(knitr)      # For dynamic report generation
```

#### 2. EVA Background

This section focuses on the data acquired using the EVA background.

##### 2.1. Data Loading and Preparation

We start by loading the dataset, filtering for the “EVA” background, and structuring the data into metadata and a spectral matrix.

Code

```
# Load the full dataset from the Excel file
# Note: For reproducibility, it's best to place the data file in the same directory as this script.
full_data <- read_excel("estudofundos.xlsx", col_names = TRUE)

# Filter the data to retain only the records for the EVA background
eva_data <- full_data %>%
  dplyr::filter(Fundo == "EVA")

# Separate metadata from the spectral data
# Here we assume spectral data starts from column 10 to 134
metadata <- eva_data %>%
  dplyr::select(1:9) # Selects all metadata columns

spectra <- as.matrix(eva_data[, 10:134]) # Creates a matrix of spectral values

# Define the wavelength range, which is necessary for some pre-processing functions like Detrend
# These values should correspond to the start and end wavenumbers of your spectra
wavenumbers <- seq(from = 901.16123805891, to = 1701.12309795625, length.out = ncol(spectra))
```

##### 2.2. Model Training Pipeline Setup

Here, we define the parameters and objects required for our automated model training and evaluation pipeline. We will test a combination of seven pre-processing techniques and four machine learning models.

Code

```
# Define the target variable (the species we want to predict)
species_labels <- as.factor(metadata$Especie)

# --- Define Pre-processing and Model Options ---
# A vector of pre-processing methods to be tested
preprocessing_options <- c("None", "Savitzky-Golay", "MSC", "SNV", "First_Derivative", "Second_Derivative", "Detrend")

# A vector of machine learning model methods available in the 'caret' package
model_options <- c("pls", "svmRadial", "knn", "rf") # PLS-DA, SVM, k-NN, Random Forest

# --- Configure Cross-Validation ---
# We use repeated 10-fold cross-validation to ensure robust performance estimates
# This process repeats the 10-fold CV 5 times with different splits
train_control <- trainControl(
  method = "repeatedcv",
  number = 10,       # Number of folds
  repeats = 5,       # Number of repetitions
  summaryFunction = defaultSummary,
  classProbs = TRUE, # Necessary for some models and metrics
  savePredictions = "final"
)

# --- Initialize Storage for Results ---
# A list to store the full trained model objects ('fit' objects) for later detailed analysis
trained_models_list <- list()

# A dataframe to store a summary of the performance metrics for quick comparison
results_summary <- data.frame(
  Preprocessing = character(),
  Model = character(),
  Accuracy_Mean = numeric(),
  Kappa_Mean = numeric(),
  stringsAsFactors = FALSE
)
```

##### 2.3. Executing the Training and Evaluation Loop

This code block iterates through each pre-processing technique and each model, applies the transformation, trains the model using cross-validation, and stores the results.

Code

```
# Loop through each pre-processing option
for (pp_method in preprocessing_options) {
  
  cat(paste0("\nApplying Pre-processing: ", pp_method, "\n"))
  
  # Apply the selected pre-processing method
  # The 'switch' statement is a clean way to handle multiple 'if/else' conditions
  processed_spectra <- switch(
    pp_method,
    "None" = spectra,
    "Savitzky-Golay" = savitzkyGolay(spectra, m = 0, p = 2, w = 15),
    "MSC" = msc(spectra),
    "SNV" = standardNormalVariate(spectra),
    "First_Derivative" = savitzkyGolay(spectra, m = 1, p = 2, w = 15),
    "Second_Derivative" = savitzkyGolay(spectra, m = 2, p = 2, w = 15),
    "Detrend" = detrend(spectra, wav = wavenumbers)
  )
  
  # Combine pre-processed spectra with species labels for training
  training_data <- as.data.frame(processed_spectra)
  training_data$species <- species_labels
  
  # Loop through each machine learning model
  for (model_method in model_options) {
    
    cat(paste0("  -> Training Model: ", model_method, "\n"))
    
    # Set a seed for reproducibility of the random processes in model training
    set.seed(123)
    
    # Use tryCatch to handle potential errors during model training without stopping the loop
    tryCatch({
      
      # Train the model using the 'train' function from the caret package
      fit <- train(
        species ~ .,
        data = training_data,
        method = model_method,
        trControl = train_control,
        metric = "Accuracy",
        # This standardizes the data (mean=0, sd=1) before training,
        # which is crucial for distance-based models like k-NN.
        preProcess = c("center", "scale")
          )
      
      # --- Store the Results ---
      
      # 1. Store the full trained model object in the list
      combination_name <- paste0(pp_method, "_", model_method)
      trained_models_list[[combination_name]] <- fit
      
      # 2. Add the key performance metrics to the summary dataframe
      results_summary <- results_summary %>%
        add_row(
          Preprocessing = pp_method,
          Model = model_method,
          Accuracy_Mean = max(fit$results$Accuracy),
          Kappa_Mean = max(fit$results$Kappa)
        )
      
    }, error = function(e) {
      # If an error occurs, print a message and add NA values to the results
      message(paste("Error training model", model_method, "with", pp_method, ":", e$message))
      results_summary <- results_summary %>%
        add_row(
          Preprocessing = pp_method,
          Model = model_method,
          Accuracy_Mean = NA,
          Kappa_Mean = NA
        )
    })
  }
}
```

```
Applying Pre-processing: None
  -> Training Model: pls
  -> Training Model: svmRadial
  -> Training Model: knn
  -> Training Model: rf

Applying Pre-processing: Savitzky-Golay
  -> Training Model: pls
  -> Training Model: svmRadial
  -> Training Model: knn
  -> Training Model: rf

Applying Pre-processing: MSC
  -> Training Model: pls
  -> Training Model: svmRadial
  -> Training Model: knn
  -> Training Model: rf

Applying Pre-processing: SNV
  -> Training Model: pls
  -> Training Model: svmRadial
  -> Training Model: knn
  -> Training Model: rf

Applying Pre-processing: First_Derivative
  -> Training Model: pls
  -> Training Model: svmRadial
  -> Training Model: knn
  -> Training Model: rf

Applying Pre-processing: Second_Derivative
  -> Training Model: pls
  -> Training Model: svmRadial
  -> Training Model: knn
  -> Training Model: rf

Applying Pre-processing: Detrend
  -> Training Model: pls
```

```
  -> Training Model: svmRadial
  -> Training Model: knn
  -> Training Model: rf
```

Code

```
cat("\n--- Model training and evaluation complete. ---\n")
```

```
--- Model training and evaluation complete. ---
```

##### 2.4. Analysis and Visualization of Results

After the training loop is complete, we can analyze and visualize the performance of all combinations.

###### 2.4.1. Ranked Performance Table

We first print a table of the results, ordered from the highest to the lowest mean accuracy, to identify the top-performing combinations.

Code

```
# Order the results by mean accuracy in descending order
ranked_results <- results_summary %>%
  arrange(desc(Accuracy_Mean))

# Print the ranked table
kable(ranked_results, caption = "Ranked Performance of Model-Preprocessing Combinations on EVA Background.")
```

Ranked Performance of Model-Preprocessing Combinations on EVA Background.

| Preprocessing | Model | Accuracy\_Mean | Kappa\_Mean |
| --- | --- | --- | --- |
| First\_Derivative | knn | 0.9879762 | 0.9836384 |
| Second\_Derivative | knn | 0.9836190 | 0.9771402 |
| Second\_Derivative | rf | 0.9725476 | 0.9621402 |
| First\_Derivative | rf | 0.9615952 | 0.9482804 |
| Detrend | rf | 0.9520714 | 0.9354645 |
| MSC | rf | 0.9344524 | 0.9116807 |
| SNV | knn | 0.9293333 | 0.9043094 |
| Detrend | knn | 0.9282619 | 0.9023923 |
| MSC | knn | 0.9227857 | 0.8956789 |
| Second\_Derivative | svmRadial | 0.9226667 | 0.8961485 |
| First\_Derivative | svmRadial | 0.9182619 | 0.8890267 |
| SNV | rf | 0.9115952 | 0.8804925 |
| Second\_Derivative | pls | 0.8800000 | 0.8363637 |
| None | svmRadial | 0.8571667 | 0.8058089 |
| Savitzky-Golay | svmRadial | 0.8543095 | 0.8019200 |
| First\_Derivative | pls | 0.8372619 | 0.7797984 |
| Detrend | svmRadial | 0.8182143 | 0.7557181 |
| None | rf | 0.8122381 | 0.7451011 |
| Savitzky-Golay | rf | 0.8045238 | 0.7350126 |
| MSC | svmRadial | 0.8016429 | 0.7352689 |
| SNV | svmRadial | 0.7904048 | 0.7193245 |
| SNV | pls | 0.7718333 | 0.6947280 |
| None | pls | 0.7655000 | 0.6841194 |
| Savitzky-Golay | pls | 0.7559762 | 0.6714971 |
| Detrend | pls | 0.7423571 | 0.6529581 |
| Savitzky-Golay | knn | 0.7141429 | 0.6165789 |
| None | knn | 0.7134286 | 0.6156780 |
| MSC | pls | 0.6346905 | 0.5121481 |

###### 2.4.2. Graphical Performance Comparison

Bar charts are used to visually compare the performance (Accuracy and Kappa) across different models and pre-processing techniques.

###### 2.4.2.1 Accuracy Plot Code

This chunk first prepares the data by calculating the overall average accuracy for each pre-processing method and then uses that average to reorder the x-axis. The plot is then built with layers for the individual model bars and the overall average line/points.

Code

```
# --- 1. Data Preparation (com um passo extra) ---
library(dplyr)
library(forcats)
library(ggplot2)

# Adicionamos o cálculo da altura máxima de cada grupo
plot_data_acc <- ranked_results %>%
  group_by(Preprocessing) %>%
  mutate(
    Avg_Accuracy = mean(Accuracy_Mean, na.rm = TRUE),
    Max_Height = max(Accuracy_Mean, na.rm = TRUE) # Altura da barra mais alta no grupo
  ) %>%
  ungroup() %>%
  mutate(Preprocessing = fct_reorder(Preprocessing, Avg_Accuracy, .desc = TRUE))

# Dataframe resumido para os marcadores da média
avg_summary_data <- plot_data_acc %>%
  distinct(Preprocessing, Avg_Accuracy, Max_Height)

# --- 2. Plot Generation (Alternativa Recomendada) ---

# Definimos um pequeno espaço vertical para o marcador da média
vertical_offset <- 0.03 

ggplot(data = plot_data_acc, aes(x = Preprocessing)) +
  
  # Camada 1 e 2: Barras e seus rótulos (sem alteração)
  geom_bar(
    aes(y = Accuracy_Mean, fill = Model),
    stat = "identity", 
    position = position_dodge(width = 0.9)
  ) +
  geom_text(
    aes(y = Accuracy_Mean, group = Model, label = round(Accuracy_Mean, 2)),
    position = position_dodge(width = 0.9),
    vjust = -0.3, 
    size = 3
  ) +
  
  # --- NOVA ABORDAGEM PARA A MÉDIA ---
  
  # Camada 3: Ponto para a média, posicionado ACIMA da barra mais alta
  geom_point(
    data = avg_summary_data,
    aes(y = Max_Height + vertical_offset), # Posição vertical dinâmica
    color = "#565656",
    size = 4,
    shape = 23, # Diamante com preenchimento
    fill = "grey"
  ) +
  
  # Camada 4: Rótulo para o ponto da média
  geom_text(
    data = avg_summary_data,
    aes(y = Max_Height + vertical_offset, label = round(Avg_Accuracy, 2)),
    vjust = -1,
    color = "#565656",
    fontface = "bold",
    size = 3.5
  ) +
  
  # Labels and Theme (com subtítulo atualizado)
  labs(
    title = "Mean Accuracy by Pre-processing and Model",
    subtitle = "Bars show individual models. The diamond marks the average for each pre-processing method.",
    x = "Pre-processing Method",
    y = "Mean Accuracy (10-fold CV, 5 repeats)"
  ) +
  # Aumenta um pouco o limite do eixo para dar espaço ao novo marcador
  scale_y_continuous(limits = c(0, 1.1), breaks = seq(0, 1, 0.2)) +
  theme_bw(base_size = 12) +
  theme(
    axis.text.x = element_text(angle = 45, hjust = 1),
    panel.grid.major.x = element_blank(),
    panel.grid.minor = element_blank()
  )
```

###### 2.4.2.2 Kappa Plot Code

The same logic is applied here for the Kappa metric. The data is prepared to reorder the x-axis based on the average Kappa score per pre-processing method, and the plot is generated accordingly.

Code

```
# --- 1. Data Preparation (com o cálculo da altura máxima) ---
library(dplyr)
library(forcats)
library(ggplot2)

# Adicionamos o cálculo da altura máxima de cada grupo para o Kappa
plot_data_kappa <- ranked_results %>%
  group_by(Preprocessing) %>%
  mutate(
    Avg_Kappa = mean(Kappa_Mean, na.rm = TRUE),
    Max_Height = max(Kappa_Mean, na.rm = TRUE) # Altura da barra mais alta no grupo
  ) %>%
  ungroup() %>%
  mutate(Preprocessing = fct_reorder(Preprocessing, Avg_Kappa, .desc = TRUE))

# Dataframe resumido para os marcadores da média
avg_summary_data_kappa <- plot_data_kappa %>%
  distinct(Preprocessing, Avg_Kappa, Max_Height)


# --- 2. Plot Generation (Alternativa Recomendada para o Kappa) ---

# Definimos um pequeno espaço vertical para o marcador da média
vertical_offset <- 0.03 

ggplot(data = plot_data_kappa, aes(x = Preprocessing)) +
  
  # Camada 1 e 2: Barras e seus rótulos (sem alteração)
  geom_bar(
    aes(y = Kappa_Mean, fill = Model),
    stat = "identity", 
    position = position_dodge(width = 0.9)
  ) +
  geom_text(
    aes(y = Kappa_Mean, group = Model, label = round(Kappa_Mean, 2)),
    position = position_dodge(width = 0.9),
    vjust = -0.3, 
    size = 3
  ) +
  
  # --- NOVA ABORDAGEM PARA A MÉDIA ---
  
  # Camada 3: Ponto para a média, posicionado ACIMA da barra mais alta
  geom_point(
    data = avg_summary_data_kappa,
    aes(y = Max_Height + vertical_offset), # Posição vertical dinâmica
    color = "#565656",
    size = 4,
    shape = 23, # Diamante com preenchimento
    fill = "grey"
  ) +
  
  # Camada 4: Rótulo para o ponto da média
  geom_text(
    data = avg_summary_data_kappa,
    aes(y = Max_Height + vertical_offset, label = round(Avg_Kappa, 2)),
    vjust = -1,
    color = "#565656",
    fontface = "bold",
    size = 3.5
  ) +

  # Labels and Theme (com subtítulo e limite do eixo atualizados)
  labs(
    title = "Mean Kappa by Pre-processing and Model",
    subtitle = "Bars show individual models. The diamond marks the average for each pre-processing method.",
    x = "Pre-processing Method",
    y = "Mean Kappa (10-fold CV, 5 repeats)"
  ) +
  # Aumenta um pouco o limite do eixo para dar espaço ao novo marcador
  scale_y_continuous(limits = c(0, 1.1), breaks = seq(0, 1, 0.2)) +
  theme_bw(base_size = 12) +
  theme(
    axis.text.x = element_text(angle = 45, hjust = 1),
    panel.grid.major.x = element_blank(),
    panel.grid.minor = element_blank()
  )
```

##### **2.5 Comprehensive Pipeline for Pre-processing and Model Evaluation**

To systematically evaluate the impact of different spectral pre-processing techniques and classification algorithms, a comprehensive and automated pipeline was developed in R. This approach ensures that every combination of pre-processing and modeling is tested under the same cross-validation scheme, allowing for a robust and unbiased comparison of performance.

The pipeline is structured in three logical steps: setup, batch pre-processing, and automated model training.

###### **2.5.1 Pipeline Setup and Configuration**

The first step involves defining all parameters and creating the necessary data structures for the analysis. This includes specifying the list of pre-processing techniques and the classification models to be evaluated. The selection of models includes a standard chemometric algorithm (PLS-DA), a distance-based method (k-NN), and widely used machine learning algorithms (SVM, Random Forest) to thoroughly explore the predictive potential of the spectral data.

Code

```
# --- Setup: Define options and create empty objects to store results ---

# A named list to store each version of the pre-processed spectra
preprocessed_spectra_list <- list()

# A named list to store all the trained 'fit' objects from caret
trained_models_list <- list()

# A dataframe to store the summary of results
results_summary <- data.frame()

# Define the preprocessing methods to be tested
preprocessing_options <- c("None", "Savitzky-Golay", "MSC", "SNV", 
                         "First_Derivative", "Second_Derivative", "Detrend")

# Define the machine learning models to be tested
model_options <- c("pls", "svmRadial", "knn", "rf")

# Define the cross-validation control parameters
train_control <- trainControl(
  method = "repeatedcv",
  number = 10,
  repeats = 5,
  summaryFunction = defaultSummary,
  classProbs = TRUE,
  savePredictions = "final"
)
```

###### **2.5.2 Stage 1: Batch Spectral Pre-processing**

The first computational stage of the pipeline is dedicated exclusively to pre-processing. The code iterates through each selected technique, applies it to the original raw spectral matrix (`spectra`), and stores the resulting transformed matrix in a named list. The original, unprocessed spectra are also included under the name `"None"` to serve as a performance baseline. This batch-processing approach is highly efficient, as it avoids redundant calculations during the subsequent modeling stage.

Code

```
# --- STAGE 1: Pre-processing Loop ---
# This loop's only job is to apply and store the results of each pre-processing.

cat("--- STAGE 1: Applying all pre-processing methods ---\n")
```

```
--- STAGE 1: Applying all pre-processing methods ---
```

Code

```
# Always include the original data as a baseline
preprocessed_spectra_list[["None"]] <- spectra

# Loop over options, apply the corresponding function, and store the result
for (pp_method in preprocessing_options) {
  # Skip "None" as it's already added
  if (pp_method == "None") next
  
  cat("  -> Applying and storing:", pp_method, "\n")
  
  processed_spectra <- switch(
    pp_method,
    "Savitzky-Golay"    = savitzkyGolay(spectra, m = 0, p = 2, w = 15),
    "MSC"               = msc(spectra),
    "SNV"               = standardNormalVariate(spectra),
    "First_Derivative"  = savitzkyGolay(spectra, m = 1, p = 2, w = 15),
    "Second_Derivative" = savitzkyGolay(spectra, m = 2, p = 2, w = 15),
    "Detrend"           = detrend(spectra, wav = wavenumbers)
  )
  
  preprocessed_spectra_list[[pp_method]] <- processed_spectra
}
```

```
  -> Applying and storing: Savitzky-Golay 
  -> Applying and storing: MSC 
  -> Applying and storing: SNV 
  -> Applying and storing: First_Derivative 
  -> Applying and storing: Second_Derivative 
  -> Applying and storing: Detrend
```

Code

```
cat("--- STAGE 1 COMPLETE: All spectra versions are stored. ---\n\n")
```

```
--- STAGE 1 COMPLETE: All spectra versions are stored. ---
```

###### **2.5.3 Stage 2: Automated Model Training and Evaluation**

The second stage of the pipeline performs the model training and evaluation. It systematically iterates through each pre-processed dataset created in Stage 1. For each dataset, an inner loop trains and evaluates every classification model defined in the setup.

The `caret::train` function is used to handle the entire process, including the application of data scaling (`preProcess = c("center", "scale")`) which is critical for distance-based algorithms like k-NN. A `tryCatch` block is implemented to ensure that the pipeline continues to run even if a specific combination of pre-processing and modeling fails. The complete trained model object and a summary of its performance metrics (Mean Accuracy and Mean Kappa) are stored for the final analysis presented in the main paper.

Code

```
# --- STAGE 2: Model Training Loop ---
# This loop iterates through the LIST of pre-processed data we just created.

cat("--- STAGE 2: Training models on each pre-processed dataset ---\n")
```

```
--- STAGE 2: Training models on each pre-processed dataset ---
```

Code

```
for (pp_name in names(preprocessed_spectra_list)) {
  
  cat(paste0("\n--- Using dataset: ", pp_name, " ---\n"))
  
  # Create the training dataframe for this iteration
  current_spectra <- preprocessed_spectra_list[[pp_name]]
  training_data <- as.data.frame(current_spectra)
  training_data$species <- species_labels
  
  # Inner loop for the models
  for (model_method in model_options) {
    cat(paste0("  -> Training model: ", model_method, "...\n"))
    
    set.seed(123) # for reproducibility
    
    tryCatch({
      fit <- train(
        species ~ .,
        data = training_data,
        method = model_method,
        trControl = train_control,
        metric = "Accuracy",
        # Standardize data (center/scale) before training
        preProcess = c("center", "scale")
      )
      
      # Store the complete trained model object
      combination_name <- paste0(pp_name, "_", model_method)
      trained_models_list[[combination_name]] <- fit
      
      # Add the key performance metrics to the summary dataframe
      results_summary <- dplyr::bind_rows(
        results_summary,
        data.frame(
          Preprocessing = pp_name,
          Model = model_method,
          Accuracy_Mean = max(fit$results$Accuracy),
          Kappa_Mean = max(fit$results$Kappa)
        )
      )
      
    }, error = function(e) {
      message(paste("ERROR training model", model_method, "with", pp_name, ":", e$message))
    })
  }
}
```

```
--- Using dataset: None ---
  -> Training model: pls...
  -> Training model: svmRadial...
  -> Training model: knn...
  -> Training model: rf...

--- Using dataset: Savitzky-Golay ---
  -> Training model: pls...
  -> Training model: svmRadial...
  -> Training model: knn...
  -> Training model: rf...

--- Using dataset: MSC ---
  -> Training model: pls...
  -> Training model: svmRadial...
  -> Training model: knn...
  -> Training model: rf...

--- Using dataset: SNV ---
  -> Training model: pls...
  -> Training model: svmRadial...
  -> Training model: knn...
  -> Training model: rf...

--- Using dataset: First_Derivative ---
  -> Training model: pls...
  -> Training model: svmRadial...
  -> Training model: knn...
  -> Training model: rf...

--- Using dataset: Second_Derivative ---
  -> Training model: pls...
  -> Training model: svmRadial...
  -> Training model: knn...
  -> Training model: rf...

--- Using dataset: Detrend ---
  -> Training model: pls...
```

```
  -> Training model: svmRadial...
  -> Training model: knn...
  -> Training model: rf...
```

Code

```
cat("\n--- STAGE 2 COMPLETE: All models have been trained. ---\n")
```

```
--- STAGE 2 COMPLETE: All models have been trained. ---
```

##### **2.6 Graphical Analysis of Cross-Validation Confusion Matrices**

To move beyond single performance metrics like Accuracy and Kappa, a detailed analysis of the confusion matrices from the cross-validation predictions was performed. A confusion matrix provides deeper insight into a model’s behavior, revealing not just its overall performance, but also its specific error patterns (e.g., which species are frequently confused with one another).

For a clear and comparative visualization, all confusion matrices were normalized by the true class (rows) to show the proportion of predictions. These normalized matrices were then plotted as a faceted heatmap. In this format, each tile represents a cell in the confusion matrix, and its color intensity corresponds to the prediction proportion. This allows for rapid visual identification of strong diagonal patterns (high classification accuracy) and specific off-diagonal hotspots (systematic misclassification errors) across all tested model and pre-processing combinations.

Code

```
# --- 1. Data Extraction and Preparation ---
# This step extracts the prediction results saved during cross-validation
# from every model stored in 'trained_models_list'.

cat("--- Extracting predictions from all trained models... ---\n")
```

```
--- Extracting predictions from all trained models... ---
```

Code

```
all_predictions <- purrr::map_dfr(
  trained_models_list,
  ~ as.data.frame(.x$pred),
  .id = "Combination"
) %>%
  # Separate the 'Combination' column into 'Preprocessing' and 'Model'
  tidyr::separate(Combination, into = c("Preprocessing", "Model"), sep = "_", extra = "merge") %>%
  # Ensure the factor levels are ordered nicely for plotting
  mutate(
    Preprocessing = fct_inorder(Preprocessing),
    Model = fct_inorder(Model)
  )

# --- 2. Calculate Normalized Confusion Matrix Proportions ---
# We group the data by each combination and true class (obs) to calculate
# the proportion of predictions made for that class.

cat("--- Calculating normalized confusion matrix proportions... ---\n")
```

```
--- Calculating normalized confusion matrix proportions... ---
```

Code

```
confusion_proportions <- all_predictions %>%
  # Count occurrences of each prediction (pred) for each true class (obs)
  group_by(Preprocessing, Model, obs, pred) %>%
  summarise(N = n(), .groups = "drop") %>%
  # Calculate the proportion based on the total for each true class
  group_by(Preprocessing, Model, obs) %>%
  mutate(Proportion = N / sum(N)) %>%
  # Ensure column names are clear for plotting
  rename(
    True_Class = obs,
    Predicted_Class = pred
  )


# --- 3. Generate the Faceted Heatmap Plot ---

cat("--- Generating the plot... ---\n")
```

```
--- Generating the plot... ---
```

Code

```
ggplot(confusion_proportions, aes(x = Predicted_Class, y = True_Class, fill = Proportion)) +
  geom_tile(color = "white", linewidth = 0.5) +
  geom_text(
    aes(label = scales::percent(Proportion, accuracy = 0.1),
        color = ifelse(Proportion > 0.5, "white", "black")),
    size = 2.5
  ) +
  
  facet_wrap(~ Preprocessing + Model, ncol = 8) +
  
  scale_fill_viridis_c(labels = scales::percent, name = "Proportion") +
  theme_bw(base_size = 10) +
  theme(
    axis.text.x = element_text(angle = 45, hjust = 1, size = 7),
    axis.text.y = element_text(size = 7),
    strip.text = element_text(face = "bold", size = 8),
    legend.position = "bottom",
    panel.grid = element_blank(),
    aspect.ratio = 1
  ) +
  labs(
    x = "Predicted Class",
    y = "True Class",
    title = "Faceted Confusion Matrices from Cross-Validation",
    subtitle = "Values and colors represent the proportion of predictions for each true class."
  ) +
  scale_color_manual(values = c("black" = "black", "white" = "white"), guide = "none")
```

##### **2.7 Visualization of Mean Spectra by Pre-processing Method**

To visually inspect the effect of each pre-processing technique on the spectral signatures of the different species, the mean spectrum for each species was calculated and plotted. This visualization allows for a qualitative assessment of how different methods enhance or alter spectral features, such as baseline shifts, peak separation, and overall spectral shape.

The following R code first transforms the list of processed spectra into a single, tidy data frame. It then calculates the mean absorbance at each wavenumber for every species-and-preprocessing combination. Finally, it generates a faceted plot, providing a direct visual comparison of the pre-processing outcomes.

Code

```
# Carregar pacotes para manipulação de gráficos
library(patchwork)
library(cowplot)
library(ggplot2)

# --- 1. Data Restructuring and 2. Calculate Mean Spectra ---

cat("--- Restructuring data for plotting... ---\n")
```

```
--- Restructuring data for plotting... ---
```

Code

```
long_spectra_data <- purrr::map_dfr(
  preprocessed_spectra_list,
  ~ {
    df <- as.data.frame(.x)
    if (ncol(df) != length(wavenumbers)) {
      trim_each_side <- (length(wavenumbers) - ncol(df)) / 2
      wavenumbers_trimmed <- wavenumbers[(trim_each_side + 1):(length(wavenumbers) - trim_each_side)]
      colnames(df) <- wavenumbers_trimmed
    } else {
      colnames(df) <- wavenumbers
    }
    df$Species <- species_labels
    df$Sample_ID <- 1:nrow(df)
    return(df)
  },
  .id = "Preprocessing"
) %>%
  tidyr::pivot_longer(
    cols = -c(Species, Sample_ID, Preprocessing),
    names_to = "Wavenumber",
    values_to = "Absorbance"
  ) %>%
  mutate(Wavenumber = as.numeric(Wavenumber))

cat("--- Calculating mean spectra for each group... ---\n")
```

```
--- Calculating mean spectra for each group... ---
```

Code

```
mean_spectra <- long_spectra_data %>%
  group_by(Preprocessing, Species, Wavenumber) %>%
  summarise(Mean_Absorbance = mean(Absorbance, na.rm = TRUE), .groups = "drop") %>%
  mutate(Preprocessing = fct_inorder(Preprocessing))


# --- 3. Generate and Combine Plots ---

cat("--- Generating final composite plot... ---\n")
```

```
--- Generating final composite plot... ---
```

Code

```
# Passo 3.1: Criar uma lista de gráficos individuais
plot_list <- lapply(unique(mean_spectra$Preprocessing), function(method_name) {
  p <- ggplot(mean_spectra[mean_spectra$Preprocessing == method_name, ], 
              aes(x = Wavenumber, y = Mean_Absorbance, color = Species)) +
    geom_line(linewidth = 0.8) +
    ggtitle(method_name) +
    scale_x_reverse() +
    scale_y_continuous(labels = scales::scientific) +
    scale_color_viridis_d() +
    theme_bw(base_size = 10) +
    theme(
      legend.position = "none",
      plot.title = element_text(face = "bold", size = 12),
      axis.title = element_blank() # Remove títulos dos eixos individuais
    )
  return(p)
})

# Passo 3.2: Criar a legenda de forma robusta
legend_source_plot <- ggplot(mean_spectra, aes(x = Wavenumber, y = Mean_Absorbance, color = Species)) +
  geom_line() +
  scale_color_viridis_d(name = "Species (Tombo)") +
  theme(legend.position = "center",
        legend.box = "vertical",
        legend.title = element_text(face="bold", size = 14),
        legend.text = element_text(size = 12),
        legend.key.size = unit(1.5, "lines"))

extracted_legend <- get_legend(legend_source_plot)

# Passo 3.3: Converter o objeto da legenda em um gráfico desenhável com ggdraw()
# Adiciona um título ao painel da legenda para clareza
legend_panel <- ggdraw(extracted_legend) +
  theme(plot.background = element_rect(fill="white", color = NA))


# Passo 3.4: Adicionar o painel da legenda à lista de gráficos
all_plots <- c(plot_list, list(legend_panel))

# Passo 3.5: Montar o painel final com patchwork
final_plot <- wrap_plots(all_plots, ncol = 4) +
  plot_annotation(
    title = "Mean FTIR Spectra by Species and Pre-processing Method",
    subtitle = "Each panel shows the mean spectra after applying the specified data treatment.",
    # Adicionar os rótulos dos eixos aqui, no tema geral da anotação
    theme = theme(
      plot.title = element_text(face = "bold", size = 18, hjust = 0.5),
      plot.subtitle = element_text(hjust = 0.5)
    )
  )

# Adiciona os eixos comuns usando o operador '&' do patchwork
final_plot <- final_plot & 
  theme(
    axis.title.x = element_text(size = 12),
    axis.title.y = element_text(size = 12)
  ) &
  labs(
    x = bquote("Wavenumber" ~ (cm^-1)),
    y = "Absorbance (or derivative)"
  )


# Exibe o gráfico final
final_plot
```

### 3. Shiny Alluminiun Background


###### Source Code

```
---
title: "Supplementary Material: Analysis of FTIR Spectra on Different Backgrounds"
format:
  html:
    toc: true
    toc-location: left
    code-fold: true
    code-tools: true
editor: visual
---

# Introduction

This document provides the supplementary material for our study on plant species identification using Fourier-transform infrared (FTIR) spectroscopy. Here, we detail the computational methodology used to test and compare the performance of various machine learning models combined with different spectral pre-processing techniques.

# 1. Setup: Loading Required Packages

First, we load all the necessary R packages for data manipulation, pre-processing, modeling, and visualization. A brief description of each package's role is provided in the comments.

```{r setup-packages, message=FALSE, warning=FALSE}

### Data Manipulation and Visualization
library(readxl)     # To read data from Excel files
library(tidyverse)  # A collection of R packages for data science (incl. dplyr, ggplot2)
library(ggplot2)    # For creating elegant and complex plots
library(ggpubr)     # For arranging and customizing ggplot2 plots
library(viridis)    # For colorblind-friendly color palettes

### Spectral Data Pre-processing
library(prospectr)  # Provides a wide range of chemometric tools, including pre-processing
library(signal)     # Used for signal processing functions like Savitzky-Golay filter

### Machine Learning and Model Evaluation
library(caret)      # Comprehensive framework for training and evaluating machine learning models
library(pls)        # Implements Partial Least Squares (used for PLS-DA)
library(randomForest) # Implements Random Forest algorithm
library(kernlab)    # Implements Support Vector Machines
library(e1071)      # General-purpose package with various ML functions

### Reporting
library(knitr)      # For dynamic report generation
```

## 2. EVA Background

This section focuses on the data acquired using the EVA background.

### 2.1. Data Loading and Preparation

We start by loading the dataset, filtering for the "EVA" background, and structuring the data into metadata and a spectral matrix.

```{r}
### Load the full dataset from the Excel file
### Note: For reproducibility, it's best to place the data file in the same directory as this script.
full_data <- read_excel("estudofundos.xlsx", col_names = TRUE)

### Filter the data to retain only the records for the EVA background
eva_data <- full_data %>%
  dplyr::filter(Fundo == "EVA")

### Separate metadata from the spectral data
### Here we assume spectral data starts from column 10 to 134
metadata <- eva_data %>%
  dplyr::select(1:9) # Selects all metadata columns

spectra <- as.matrix(eva_data[, 10:134]) # Creates a matrix of spectral values

### Define the wavelength range, which is necessary for some pre-processing functions like Detrend
### These values should correspond to the start and end wavenumbers of your spectra
wavenumbers <- seq(from = 901.16123805891, to = 1701.12309795625, length.out = ncol(spectra))
```

### 2.2. Model Training Pipeline Setup

Here, we define the parameters and objects required for our automated model training and evaluation pipeline. We will test a combination of seven pre-processing techniques and four machine learning models.

```{r}
### Define the target variable (the species we want to predict)
species_labels <- as.factor(metadata$Especie)

### --- Define Pre-processing and Model Options ---
### A vector of pre-processing methods to be tested
preprocessing_options <- c("None", "Savitzky-Golay", "MSC", "SNV", "First_Derivative", "Second_Derivative", "Detrend")

### A vector of machine learning model methods available in the 'caret' package
model_options <- c("pls", "svmRadial", "knn", "rf") # PLS-DA, SVM, k-NN, Random Forest

### --- Configure Cross-Validation ---
### We use repeated 10-fold cross-validation to ensure robust performance estimates
### This process repeats the 10-fold CV 5 times with different splits
train_control <- trainControl(
  method = "repeatedcv",
  number = 10,       # Number of folds
  repeats = 5,       # Number of repetitions
  summaryFunction = defaultSummary,
  classProbs = TRUE, # Necessary for some models and metrics
  savePredictions = "final"
)

### --- Initialize Storage for Results ---
### A list to store the full trained model objects ('fit' objects) for later detailed analysis
trained_models_list <- list()

### A dataframe to store a summary of the performance metrics for quick comparison
results_summary <- data.frame(
  Preprocessing = character(),
  Model = character(),
  Accuracy_Mean = numeric(),
  Kappa_Mean = numeric(),
  stringsAsFactors = FALSE
)
```

### 2.3. Executing the Training and Evaluation Loop

This code block iterates through each pre-processing technique and each model, applies the transformation, trains the model using cross-validation, and stores the results.

```{r}
#| warning: false
#| 
### Loop through each pre-processing option
for (pp_method in preprocessing_options) {
  
  cat(paste0("\nApplying Pre-processing: ", pp_method, "\n"))
  
  # Apply the selected pre-processing method
  # The 'switch' statement is a clean way to handle multiple 'if/else' conditions
  processed_spectra <- switch(
    pp_method,
    "None" = spectra,
    "Savitzky-Golay" = savitzkyGolay(spectra, m = 0, p = 2, w = 15),
    "MSC" = msc(spectra),
    "SNV" = standardNormalVariate(spectra),
    "First_Derivative" = savitzkyGolay(spectra, m = 1, p = 2, w = 15),
    "Second_Derivative" = savitzkyGolay(spectra, m = 2, p = 2, w = 15),
    "Detrend" = detrend(spectra, wav = wavenumbers)
  )
  
  # Combine pre-processed spectra with species labels for training
  training_data <- as.data.frame(processed_spectra)
  training_data$species <- species_labels
  
  # Loop through each machine learning model
  for (model_method in model_options) {
    
    cat(paste0("  -> Training Model: ", model_method, "\n"))
    
    # Set a seed for reproducibility of the random processes in model training
    set.seed(123)
    
    # Use tryCatch to handle potential errors during model training without stopping the loop
    tryCatch({
      
      # Train the model using the 'train' function from the caret package
      fit <- train(
        species ~ .,
        data = training_data,
        method = model_method,
        trControl = train_control,
        metric = "Accuracy",
        # This standardizes the data (mean=0, sd=1) before training,
        # which is crucial for distance-based models like k-NN.
        preProcess = c("center", "scale")
          )
      
      # --- Store the Results ---
      
      # 1. Store the full trained model object in the list
      combination_name <- paste0(pp_method, "_", model_method)
      trained_models_list[[combination_name]] <- fit
      
      # 2. Add the key performance metrics to the summary dataframe
      results_summary <- results_summary %>%
        add_row(
          Preprocessing = pp_method,
          Model = model_method,
          Accuracy_Mean = max(fit$results$Accuracy),
          Kappa_Mean = max(fit$results$Kappa)
        )
      
    }, error = function(e) {
      # If an error occurs, print a message and add NA values to the results
      message(paste("Error training model", model_method, "with", pp_method, ":", e$message))
      results_summary <- results_summary %>%
        add_row(
          Preprocessing = pp_method,
          Model = model_method,
          Accuracy_Mean = NA,
          Kappa_Mean = NA
        )
    })
  }
}

cat("\n--- Model training and evaluation complete. ---\n")
```

### 2.4. Analysis and Visualization of Results

After the training loop is complete, we can analyze and visualize the performance of all combinations.

#### 2.4.1. Ranked Performance Table

We first print a table of the results, ordered from the highest to the lowest mean accuracy, to identify the top-performing combinations.

```{r}
### Order the results by mean accuracy in descending order
ranked_results <- results_summary %>%
  arrange(desc(Accuracy_Mean))

### Print the ranked table
kable(ranked_results, caption = "Ranked Performance of Model-Preprocessing Combinations on EVA Background.")
```

#### 2.4.2. Graphical Performance Comparison

Bar charts are used to visually compare the performance (Accuracy and Kappa) across different models and pre-processing techniques.

#### 2.4.2.1 Accuracy Plot Code

This chunk first prepares the data by calculating the overall average accuracy for each pre-processing method and then uses that average to reorder the x-axis. The plot is then built with layers for the individual model bars and the overall average line/points.

```{r}
### --- 1. Data Preparation (com um passo extra) ---
library(dplyr)
library(forcats)
library(ggplot2)

### Adicionamos o cálculo da altura máxima de cada grupo
plot_data_acc <- ranked_results %>%
  group_by(Preprocessing) %>%
  mutate(
    Avg_Accuracy = mean(Accuracy_Mean, na.rm = TRUE),
    Max_Height = max(Accuracy_Mean, na.rm = TRUE) # Altura da barra mais alta no grupo
  ) %>%
  ungroup() %>%
  mutate(Preprocessing = fct_reorder(Preprocessing, Avg_Accuracy, .desc = TRUE))

### Dataframe resumido para os marcadores da média
avg_summary_data <- plot_data_acc %>%
  distinct(Preprocessing, Avg_Accuracy, Max_Height)

### --- 2. Plot Generation (Alternativa Recomendada) ---

### Definimos um pequeno espaço vertical para o marcador da média
vertical_offset <- 0.03 

ggplot(data = plot_data_acc, aes(x = Preprocessing)) +
  
  # Camada 1 e 2: Barras e seus rótulos (sem alteração)
  geom_bar(
    aes(y = Accuracy_Mean, fill = Model),
    stat = "identity", 
    position = position_dodge(width = 0.9)
  ) +
  geom_text(
    aes(y = Accuracy_Mean, group = Model, label = round(Accuracy_Mean, 2)),
    position = position_dodge(width = 0.9),
    vjust = -0.3, 
    size = 3
  ) +
  
  # --- NOVA ABORDAGEM PARA A MÉDIA ---
  
  # Camada 3: Ponto para a média, posicionado ACIMA da barra mais alta
  geom_point(
    data = avg_summary_data,
    aes(y = Max_Height + vertical_offset), # Posição vertical dinâmica
    color = "#565656",
    size = 4,
    shape = 23, # Diamante com preenchimento
    fill = "grey"
  ) +
  
  # Camada 4: Rótulo para o ponto da média
  geom_text(
    data = avg_summary_data,
    aes(y = Max_Height + vertical_offset, label = round(Avg_Accuracy, 2)),
    vjust = -1,
    color = "#565656",
    fontface = "bold",
    size = 3.5
  ) +
  
  # Labels and Theme (com subtítulo atualizado)
  labs(
    title = "Mean Accuracy by Pre-processing and Model",
    subtitle = "Bars show individual models. The diamond marks the average for each pre-processing method.",
    x = "Pre-processing Method",
    y = "Mean Accuracy (10-fold CV, 5 repeats)"
  ) +
  # Aumenta um pouco o limite do eixo para dar espaço ao novo marcador
  scale_y_continuous(limits = c(0, 1.1), breaks = seq(0, 1, 0.2)) +
  theme_bw(base_size = 12) +
  theme(
    axis.text.x = element_text(angle = 45, hjust = 1),
    panel.grid.major.x = element_blank(),
    panel.grid.minor = element_blank()
  )
```

#### 2.4.2.2 Kappa Plot Code

The same logic is applied here for the Kappa metric. The data is prepared to reorder the x-axis based on the average Kappa score per pre-processing method, and the plot is generated accordingly.

```{r}
### --- 1. Data Preparation (com o cálculo da altura máxima) ---
library(dplyr)
library(forcats)
library(ggplot2)

### Adicionamos o cálculo da altura máxima de cada grupo para o Kappa
plot_data_kappa <- ranked_results %>%
  group_by(Preprocessing) %>%
  mutate(
    Avg_Kappa = mean(Kappa_Mean, na.rm = TRUE),
    Max_Height = max(Kappa_Mean, na.rm = TRUE) # Altura da barra mais alta no grupo
  ) %>%
  ungroup() %>%
  mutate(Preprocessing = fct_reorder(Preprocessing, Avg_Kappa, .desc = TRUE))

### Dataframe resumido para os marcadores da média
avg_summary_data_kappa <- plot_data_kappa %>%
  distinct(Preprocessing, Avg_Kappa, Max_Height)


### --- 2. Plot Generation (Alternativa Recomendada para o Kappa) ---

### Definimos um pequeno espaço vertical para o marcador da média
vertical_offset <- 0.03 

ggplot(data = plot_data_kappa, aes(x = Preprocessing)) +
  
  # Camada 1 e 2: Barras e seus rótulos (sem alteração)
  geom_bar(
    aes(y = Kappa_Mean, fill = Model),
    stat = "identity", 
    position = position_dodge(width = 0.9)
  ) +
  geom_text(
    aes(y = Kappa_Mean, group = Model, label = round(Kappa_Mean, 2)),
    position = position_dodge(width = 0.9),
    vjust = -0.3, 
    size = 3
  ) +
  
  # --- NOVA ABORDAGEM PARA A MÉDIA ---
  
  # Camada 3: Ponto para a média, posicionado ACIMA da barra mais alta
  geom_point(
    data = avg_summary_data_kappa,
    aes(y = Max_Height + vertical_offset), # Posição vertical dinâmica
    color = "#565656",
    size = 4,
    shape = 23, # Diamante com preenchimento
    fill = "grey"
  ) +
  
  # Camada 4: Rótulo para o ponto da média
  geom_text(
    data = avg_summary_data_kappa,
    aes(y = Max_Height + vertical_offset, label = round(Avg_Kappa, 2)),
    vjust = -1,
    color = "#565656",
    fontface = "bold",
    size = 3.5
  ) +

  # Labels and Theme (com subtítulo e limite do eixo atualizados)
  labs(
    title = "Mean Kappa by Pre-processing and Model",
    subtitle = "Bars show individual models. The diamond marks the average for each pre-processing method.",
    x = "Pre-processing Method",
    y = "Mean Kappa (10-fold CV, 5 repeats)"
  ) +
  # Aumenta um pouco o limite do eixo para dar espaço ao novo marcador
  scale_y_continuous(limits = c(0, 1.1), breaks = seq(0, 1, 0.2)) +
  theme_bw(base_size = 12) +
  theme(
    axis.text.x = element_text(angle = 45, hjust = 1),
    panel.grid.major.x = element_blank(),
    panel.grid.minor = element_blank()
  )
```

### **2.5 Comprehensive Pipeline for Pre-processing and Model Evaluation**

To systematically evaluate the impact of different spectral pre-processing techniques and classification algorithms, a comprehensive and automated pipeline was developed in R. This approach ensures that every combination of pre-processing and modeling is tested under the same cross-validation scheme, allowing for a robust and unbiased comparison of performance.

The pipeline is structured in three logical steps: setup, batch pre-processing, and automated model training.

#### **2.5.1 Pipeline Setup and Configuration**

The first step involves defining all parameters and creating the necessary data structures for the analysis. This includes specifying the list of pre-processing techniques and the classification models to be evaluated. The selection of models includes a standard chemometric algorithm (PLS-DA), a distance-based method (k-NN), and widely used machine learning algorithms (SVM, Random Forest) to thoroughly explore the predictive potential of the spectral data.

```{r}
#| label: pipeline-setup
#| warning: false
#| message: false

### --- Setup: Define options and create empty objects to store results ---

### A named list to store each version of the pre-processed spectra
preprocessed_spectra_list <- list()

### A named list to store all the trained 'fit' objects from caret
trained_models_list <- list()

### A dataframe to store the summary of results
results_summary <- data.frame()

### Define the preprocessing methods to be tested
preprocessing_options <- c("None", "Savitzky-Golay", "MSC", "SNV", 
                         "First_Derivative", "Second_Derivative", "Detrend")

### Define the machine learning models to be tested
model_options <- c("pls", "svmRadial", "knn", "rf")

### Define the cross-validation control parameters
train_control <- trainControl(
  method = "repeatedcv",
  number = 10,
  repeats = 5,
  summaryFunction = defaultSummary,
  classProbs = TRUE,
  savePredictions = "final"
)
```

#### **2.5.2 Stage 1: Batch Spectral Pre-processing**

The first computational stage of the pipeline is dedicated exclusively to pre-processing. The code iterates through each selected technique, applies it to the original raw spectral matrix (`spectra`), and stores the resulting transformed matrix in a named list. The original, unprocessed spectra are also included under the name `"None"` to serve as a performance baseline. This batch-processing approach is highly efficient, as it avoids redundant calculations during the subsequent modeling stage.

```{r}
#| label: stage1-preprocessing
#| cache: true
#| warning: false
#| message: false

### --- STAGE 1: Pre-processing Loop ---
### This loop's only job is to apply and store the results of each pre-processing.

cat("--- STAGE 1: Applying all pre-processing methods ---\n")

### Always include the original data as a baseline
preprocessed_spectra_list[["None"]] <- spectra

### Loop over options, apply the corresponding function, and store the result
for (pp_method in preprocessing_options) {
  # Skip "None" as it's already added
  if (pp_method == "None") next
  
  cat("  -> Applying and storing:", pp_method, "\n")
  
  processed_spectra <- switch(
    pp_method,
    "Savitzky-Golay"    = savitzkyGolay(spectra, m = 0, p = 2, w = 15),
    "MSC"               = msc(spectra),
    "SNV"               = standardNormalVariate(spectra),
    "First_Derivative"  = savitzkyGolay(spectra, m = 1, p = 2, w = 15),
    "Second_Derivative" = savitzkyGolay(spectra, m = 2, p = 2, w = 15),
    "Detrend"           = detrend(spectra, wav = wavenumbers)
  )
  
  preprocessed_spectra_list[[pp_method]] <- processed_spectra
}
cat("--- STAGE 1 COMPLETE: All spectra versions are stored. ---\n\n")
```

#### **2.5.3 Stage 2: Automated Model Training and Evaluation**

The second stage of the pipeline performs the model training and evaluation. It systematically iterates through each pre-processed dataset created in Stage 1. For each dataset, an inner loop trains and evaluates every classification model defined in the setup.

The `caret::train` function is used to handle the entire process, including the application of data scaling (`preProcess = c("center", "scale")`) which is critical for distance-based algorithms like k-NN. A `tryCatch` block is implemented to ensure that the pipeline continues to run even if a specific combination of pre-processing and modeling fails. The complete trained model object and a summary of its performance metrics (Mean Accuracy and Mean Kappa) are stored for the final analysis presented in the main paper.

```{r}
#| label: stage2-modeling
#| cache: true
#| warning: false
#| message: false

### --- STAGE 2: Model Training Loop ---
### This loop iterates through the LIST of pre-processed data we just created.

cat("--- STAGE 2: Training models on each pre-processed dataset ---\n")

for (pp_name in names(preprocessed_spectra_list)) {
  
  cat(paste0("\n--- Using dataset: ", pp_name, " ---\n"))
  
  # Create the training dataframe for this iteration
  current_spectra <- preprocessed_spectra_list[[pp_name]]
  training_data <- as.data.frame(current_spectra)
  training_data$species <- species_labels
  
  # Inner loop for the models
  for (model_method in model_options) {
    cat(paste0("  -> Training model: ", model_method, "...\n"))
    
    set.seed(123) # for reproducibility
    
    tryCatch({
      fit <- train(
        species ~ .,
        data = training_data,
        method = model_method,
        trControl = train_control,
        metric = "Accuracy",
        # Standardize data (center/scale) before training
        preProcess = c("center", "scale")
      )
      
      # Store the complete trained model object
      combination_name <- paste0(pp_name, "_", model_method)
      trained_models_list[[combination_name]] <- fit
      
      # Add the key performance metrics to the summary dataframe
      results_summary <- dplyr::bind_rows(
        results_summary,
        data.frame(
          Preprocessing = pp_name,
          Model = model_method,
          Accuracy_Mean = max(fit$results$Accuracy),
          Kappa_Mean = max(fit$results$Kappa)
        )
      )
      
    }, error = function(e) {
      message(paste("ERROR training model", model_method, "with", pp_name, ":", e$message))
    })
  }
}
cat("\n--- STAGE 2 COMPLETE: All models have been trained. ---\n")
```

### **2.6 Graphical Analysis of Cross-Validation Confusion Matrices**

To move beyond single performance metrics like Accuracy and Kappa, a detailed analysis of the confusion matrices from the cross-validation predictions was performed. A confusion matrix provides deeper insight into a model's behavior, revealing not just its overall performance, but also its specific error patterns (e.g., which species are frequently confused with one another).

For a clear and comparative visualization, all confusion matrices were normalized by the true class (rows) to show the proportion of predictions. These normalized matrices were then plotted as a faceted heatmap. In this format, each tile represents a cell in the confusion matrix, and its color intensity corresponds to the prediction proportion. This allows for rapid visual identification of strong diagonal patterns (high classification accuracy) and specific off-diagonal hotspots (systematic misclassification errors) across all tested model and pre-processing combinations.

```{r}
#| label: plot-confusion-matrices
#| fig-width: 20
#| fig-height: 10
#| warning: false
#| message: false

### --- 1. Data Extraction and Preparation ---
### This step extracts the prediction results saved during cross-validation
### from every model stored in 'trained_models_list'.

cat("--- Extracting predictions from all trained models... ---\n")

all_predictions <- purrr::map_dfr(
  trained_models_list,
  ~ as.data.frame(.x$pred),
  .id = "Combination"
) %>%
  # Separate the 'Combination' column into 'Preprocessing' and 'Model'
  tidyr::separate(Combination, into = c("Preprocessing", "Model"), sep = "_", extra = "merge") %>%
  # Ensure the factor levels are ordered nicely for plotting
  mutate(
    Preprocessing = fct_inorder(Preprocessing),
    Model = fct_inorder(Model)
  )

### --- 2. Calculate Normalized Confusion Matrix Proportions ---
### We group the data by each combination and true class (obs) to calculate
### the proportion of predictions made for that class.

cat("--- Calculating normalized confusion matrix proportions... ---\n")

confusion_proportions <- all_predictions %>%
  # Count occurrences of each prediction (pred) for each true class (obs)
  group_by(Preprocessing, Model, obs, pred) %>%
  summarise(N = n(), .groups = "drop") %>%
  # Calculate the proportion based on the total for each true class
  group_by(Preprocessing, Model, obs) %>%
  mutate(Proportion = N / sum(N)) %>%
  # Ensure column names are clear for plotting
  rename(
    True_Class = obs,
    Predicted_Class = pred
  )


### --- 3. Generate the Faceted Heatmap Plot ---

cat("--- Generating the plot... ---\n")

ggplot(confusion_proportions, aes(x = Predicted_Class, y = True_Class, fill = Proportion)) +
  geom_tile(color = "white", linewidth = 0.5) +
  geom_text(
    aes(label = scales::percent(Proportion, accuracy = 0.1),
        color = ifelse(Proportion > 0.5, "white", "black")),
    size = 2.5
  ) +
  
  facet_wrap(~ Preprocessing + Model, ncol = 8) +
  
  scale_fill_viridis_c(labels = scales::percent, name = "Proportion") +
  theme_bw(base_size = 10) +
  theme(
    axis.text.x = element_text(angle = 45, hjust = 1, size = 7),
    axis.text.y = element_text(size = 7),
    strip.text = element_text(face = "bold", size = 8),
    legend.position = "bottom",
    panel.grid = element_blank(),
    aspect.ratio = 1
  ) +
  labs(
    x = "Predicted Class",
    y = "True Class",
    title = "Faceted Confusion Matrices from Cross-Validation",
    subtitle = "Values and colors represent the proportion of predictions for each true class."
  ) +
  scale_color_manual(values = c("black" = "black", "white" = "white"), guide = "none")
```

### **2.7 Visualization of Mean Spectra by Pre-processing Method**

To visually inspect the effect of each pre-processing technique on the spectral signatures of the different species, the mean spectrum for each species was calculated and plotted. This visualization allows for a qualitative assessment of how different methods enhance or alter spectral features, such as baseline shifts, peak separation, and overall spectral shape.

The following R code first transforms the list of processed spectra into a single, tidy data frame. It then calculates the mean absorbance at each wavenumber for every species-and-preprocessing combination. Finally, it generates a faceted plot, providing a direct visual comparison of the pre-processing outcomes.

```{r}
#| label: plot-mean-spectra
#| fig-width: 16
#| fig-height: 8
#| warning: false
#| message: false

### Carregar pacotes para manipulação de gráficos
library(patchwork)
library(cowplot)
library(ggplot2)

### --- 1. Data Restructuring and 2. Calculate Mean Spectra ---

cat("--- Restructuring data for plotting... ---\n")

long_spectra_data <- purrr::map_dfr(
  preprocessed_spectra_list,
  ~ {
    df <- as.data.frame(.x)
    if (ncol(df) != length(wavenumbers)) {
      trim_each_side <- (length(wavenumbers) - ncol(df)) / 2
      wavenumbers_trimmed <- wavenumbers[(trim_each_side + 1):(length(wavenumbers) - trim_each_side)]
      colnames(df) <- wavenumbers_trimmed
    } else {
      colnames(df) <- wavenumbers
    }
    df$Species <- species_labels
    df$Sample_ID <- 1:nrow(df)
    return(df)
  },
  .id = "Preprocessing"
) %>%
  tidyr::pivot_longer(
    cols = -c(Species, Sample_ID, Preprocessing),
    names_to = "Wavenumber",
    values_to = "Absorbance"
  ) %>%
  mutate(Wavenumber = as.numeric(Wavenumber))

cat("--- Calculating mean spectra for each group... ---\n")

mean_spectra <- long_spectra_data %>%
  group_by(Preprocessing, Species, Wavenumber) %>%
  summarise(Mean_Absorbance = mean(Absorbance, na.rm = TRUE), .groups = "drop") %>%
  mutate(Preprocessing = fct_inorder(Preprocessing))


### --- 3. Generate and Combine Plots ---

cat("--- Generating final composite plot... ---\n")

### Passo 3.1: Criar uma lista de gráficos individuais
plot_list <- lapply(unique(mean_spectra$Preprocessing), function(method_name) {
  p <- ggplot(mean_spectra[mean_spectra$Preprocessing == method_name, ], 
              aes(x = Wavenumber, y = Mean_Absorbance, color = Species)) +
    geom_line(linewidth = 0.8) +
    ggtitle(method_name) +
    scale_x_reverse() +
    scale_y_continuous(labels = scales::scientific) +
    scale_color_viridis_d() +
    theme_bw(base_size = 10) +
    theme(
      legend.position = "none",
      plot.title = element_text(face = "bold", size = 12),
      axis.title = element_blank() # Remove títulos dos eixos individuais
    )
  return(p)
})

### Passo 3.2: Criar a legenda de forma robusta
legend_source_plot <- ggplot(mean_spectra, aes(x = Wavenumber, y = Mean_Absorbance, color = Species)) +
  geom_line() +
  scale_color_viridis_d(name = "Species (Tombo)") +
  theme(legend.position = "center",
        legend.box = "vertical",
        legend.title = element_text(face="bold", size = 14),
        legend.text = element_text(size = 12),
        legend.key.size = unit(1.5, "lines"))

extracted_legend <- get_legend(legend_source_plot)

### Passo 3.3: Converter o objeto da legenda em um gráfico desenhável com ggdraw()
### Adiciona um título ao painel da legenda para clareza
legend_panel <- ggdraw(extracted_legend) +
  theme(plot.background = element_rect(fill="white", color = NA))


### Passo 3.4: Adicionar o painel da legenda à lista de gráficos
all_plots <- c(plot_list, list(legend_panel))

### Passo 3.5: Montar o painel final com patchwork
final_plot <- wrap_plots(all_plots, ncol = 4) +
  plot_annotation(
    title = "Mean FTIR Spectra by Species and Pre-processing Method",
    subtitle = "Each panel shows the mean spectra after applying the specified data treatment.",
    # Adicionar os rótulos dos eixos aqui, no tema geral da anotação
    theme = theme(
      plot.title = element_text(face = "bold", size = 18, hjust = 0.5),
      plot.subtitle = element_text(hjust = 0.5)
    )
  )

### Adiciona os eixos comuns usando o operador '&' do patchwork
final_plot <- final_plot & 
  theme(
    axis.title.x = element_text(size = 12),
    axis.title.y = element_text(size = 12)
  ) &
  labs(
    x = bquote("Wavenumber" ~ (cm^-1)),
    y = "Absorbance (or derivative)"
  )


### Exibe o gráfico final
final_plot
```

# 3. Shiny Alluminiun Background
```
